# Orthogonal targeting of KDM6A/B and HDACs mediates potent therapeutic effects in *IDH1*-mutant glioma

**DOI:** 10.1101/2020.11.26.400234

**Authors:** Alisan Kayabolen, Ebru Yilmaz, Gizem Nur Sahin, Fidan Seker-Polat, Ahmet Cingoz, Bekir Isik, Simge Acar, Hiroaki Wakimoto, Daniel P. Cahill, Ihsan Solaroglu, Adam P Cribbs, Udo Oppermann, Tugba Bagci-Onder

**Affiliations:** Brain Cancer Research and Therapy Lab, Koç University School of Medicine, Istanbul, 34450, Turkey; Koç University Research Center for Translational Medicine (KUTTAM), Koç University, Istanbul, Turkey; Department of Histology and Embryology, Koç University School of Medicine, Istanbul, 34450, Turkey; Department of Neurosurgery, Massachusetts General Hospital Cancer Center, Harvard Medical School, Boston, Massachusetts, 02114, USA; Department of Neurosurgery, Koç University School of Medicine, Istanbul, 34450, Turkey; Botnar Research Centre, Nuffield Department of Orthopedics, Rheumatology and Musculoskeletal Sciences, National Institute of Health Research Oxford Biomedical Research Unit, University of Oxford, OX3 7LD, UK; Centre for Medicines Discovery, University of Oxford, OX3 7DQ, UK; FRIAS, Freiburg Institute of Advanced Studies, University of Freiburg, 79104, Freiburg, Germany

**Keywords:** *IDH*-mutation, glioma, epigenetic, stress response, KDM6, GSK-J4, Belinostat

## Abstract

**Background:** *IDH1/2*-mutant gliomas are primary brain tumors for which curative treatments are lacking. Mutant IDH-dependent 2-hydroxyglutarate (2-HG) accumulation leads to DNA and histone hypermethylation. Based on this distinct phenotype, we interrogated epigenetic dependencies of *IDH*-mutant glioma that can be targeted therapeutically.

**Methods:** We conducted a chemical screen targeting chromatin modifiers in patient derived *IDH1*-mutant GBM cells. We investigated mechanisms of action of compound hits and their combinations through cell-based functional assays, live-cell imaging, Western blot, CRISPR knockout, RNA-seq and ChIP experiments. The therapeutic concept was validated *in vivo* using chemical inhibitors GSK-J4 and Belinostat in an orthotopic GBM model.

**Results:** We identified the H3K27me3 demethylase (KDM6) inhibitor GSK-J4 and histone deacetylase inhibitor Belinostat as potent, genotype-selective agents against *IDH1*-mutant glioma. RNA-sequencing on paired wild-type and *IDH1*^*R132H*^ cells revealed inhibition of cholesterol biosynthesis and activation of cellular stress in *IDH1*^*R132H*^ cells, which were reversible with a mutant IDH1 inhibitor. GSK-J4 caused further repression of cholesterol biosynthesis pathway genes through H3K27me3 deposition and exacerbated the ATF4-mediated integrated stress response. Belinostat inhibited anti-apoptotic pathways through activation of TGF-β signaling and induced cell cycle arrest. Together, the GSK-J4 and Belinostat combination activated *DDIT3*/CHOP-dependent apoptosis in *IDH1*-mutant cells and extended survival in an *IDH1*-mutant orthotopic model *in vivo*.

**Conclusions:** These results provide a possible therapeutic approach that exploits epigenetic vulnerabilities of *IDH*-mutant gliomas.

**Key points:**

- Combination of GSK-J4 and Belinostat selectively targets *IDH1*-mutant cells.
- GSK-J4 downregulates cholesterol biosynthesis and activates an ATF4-mediated stress response.
- Belinostat activates the TGFβ pathway, induces G2/M arrest and inhibits anti-apoptotic pathways.

**Importance of the study:** *IDH1/2* genes are frequently mutated in low grade glioma and secondary glioblastoma. These tumors exhibit a distinct epigenomic signature with increased DNA and histone methylation; therefore, identifying and exploiting their epigenetic vulnerabilities may lead to effective therapies. We discovered that targeting of KDM6A/6B together with HDACs provides a promising therapeutic approach for *IDH1*-mutant glioma.

## INTRODUCTION

Glioblastoma (GBM) is the highest grade of gliomas and among the deadliest cancers with an average survival of 14-16 months even for patients receiving surgery and chemoradiation^1^. Survival rate is higher in low-grade glioma (LGG), however most LGGs eventually progress to high-grade glioma or secondary GBM. In an integrated genomic analysis, a breakthrough discovery showed that a subset of GBM samples carried a point mutation in Isocitrate Dehydrogenase 1 (*IDH1)* gene^2^. Later, *IDH1* mutation was found in almost 70-80% of LGGs and secondary GBMs^3^. Mutations were exclusive at position 132 in *IDH1* or position 172 in *IDH2* genes, affecting the active site responsible of converting isocitrate to alpha-ketoglutarate (2-oxoglutarate, 2-OG)^4^. Mostly, with a G-to-A conversion mutation (G395A), the wild-type residue arginine is replaced with histidine (R132H) in the IDH1 enzyme.

Point mutations in *IDH1* or *IDH2* genes lead to a gain-of-function of the enzyme. Mutant IDH converts 2-OG produced by wild-type IDH, into 2-hydroxyglutarate (2-HG)^4^. 2-HG acts as an antagonist of 2-OG and inhibits activities of many 2-OG-dependent enzymes, such as TET or KDMs, which are DNA and histone demethylases, respectively^4^. Therefore, IDH mutation leads to hypermethylation in DNA and histones generating a distinct epigenetic profile called glioma CpG island methylator phenotype (G-CIMP)^5,6^. This 2-HG-mediated hypermethylation is thought to be responsible for tumorigenesis, but mutant IDH also leads to metabolic deficiencies, such as impaired TCA cycle and metabolism^7^, inhibition of mTOR and ATP synthase^8^, and downregulation of lactate dehydrogenase A (LDHA) enzyme^9^, resulting in attenuated growth of tumor cells. Recently, mutant IDH specific inhibitors emerged as a promising strategy to delay tumor growth by preventing tumorigenic effects of mutant IDH^10–12^. On the other hand, considering the metabolic deficiencies, exploitation of specific metabolic vulnerabilities of IDH mutant cells is also considered an intriguing strategy^13–15^.

In this study, we interrogated the epigenetic vulnerabilities of *IDH1*-mutant glioma cells. From a chemical screen targeting several chromatin modifiers, we established that a combination of GSK-J4, a histone demethylase inhibitor, and Belinostat, a histone deacetylase inhibitor, was highly effective at eradicating *IDH1*-mutant glioma cells. This selective vulnerability of *IDH1*-mutant cells involved the inhibition of cholesterol biosynthesis and the activation of an integrated stress response collectively leading to apoptosis, which were reversible by treatment with the mutant IDH enzyme inhibitor GSK864. Together, our findings reveal a novel combinatorial approach for *IDH1*-mutant gliomas by exploiting their epigenetic vulnerabilities.

## MATERIALS AND METHODS

### Reagents and cell lines

All reagents and cell lines used in this study are described in **Supplementary Information**.

### Cell viability and apoptosis assays

Cell viability, caspase activation assays, YO-PRO-staining, Western Blotting for cleaved PARP and Bcl-XL are described in **Supplementary Information**.

### Chemical screen with epigenetic regulator inhibitors

The epigenetic chemical probe library was constructed as described^16^. MGG119 and MGG152 spheres were dissociated into individual cells and seeded in 96-well plates as 4000 cells/well. The next day, cells were treated with the library consisting of 46 inhibitors targeting bromodomains (BRD), histone deacetylases (HDAC), histone methyltransferases (HMT), lysine demethylases (KDM), prolyl hydroxylases (PHD), methyl lysine binders, DNA methyltransferases (DNMT), poly ADP ribose polymerase (PARP), kinase inhibitors, and histone acetyltransferases (HAT) (**Supplementary Table 1**). Cell viabilities after 48h of treatment were determined via Cell Titer-Glo (CTG) Assay as described in the **Supplementary Information**. All experiments were carried out in triplicate.

### Validation of screen hits

5-azacytidine, Chaetocin, GSK-J4 and Belinostat were applied individually in a dose-dependent manner on MGG119, MGG152, Fibro1 and Fibro2 cells and cell viability was measured after 72h of treatment. All cell lines were subjected to all possible dual combinations of these inhibitors. Human astrocytes (HA) were also treated with GSK-J4 (2.5µM) and Belinostat (1µM) individually and in combination. Combination index (CI) values were calculated using CompuSyn software^17^, for 5-azacytidine and Chaetocin or GSK-J4 and Belinostat combinations in MGG119 and MGG152 cells.

### Viral packaging and transduction

Retroviral particles from pMIG Bcl-xL (Addgene #8790), and lentiviral particles from pLenti6.3/TO/V5 containing IDH1^R132H^,^13^ or pLentiCRISPRv2 (Addgene #52961) were produced in 293T cells as previously described^18^. Further details are described in **Supplementary Information**.

### Generation of *IDH1-*mutant GBM cell lines

*IDH1*-mutant cells were generated by lentiviral infection of pLenti6.3/TO/V5 containing *IDH1*^*R132H*^ cDNA^13^. Cells were cultured in parallel with wild-type cells to obtain a paired cell line. Mutant IDH1 expression was analyzed with western blot and immunocytochemical staining using IDH1^R132H^ specific antibody (DIA-H09, Dianova, Germany). 2-HG production was assessed via D-2-Hydroxyglutarate (D2HG) Assay Kit (Sigma-Aldrich, USA). Paired GBM cell lines were also treated with hit inhibitors individually and in combination. Cell viabilities were determined after 72h.

### CRISPR-mediated knock-out and activation studies

gRNA sequences targeting exon regions of KDM6A and KDM6B genes were obtained from GeCKO library designed by Feng Zhang’s laboratory at Broad Institute^19^. gRNAs targeting DDIT3 gene or LDLR promoter were designed using Chopchop gRNA design tool^20^. Details of cloning and transduction are given in **Supplementary Information**. Gene knockouts were validated via Sanger sequencing of targeted regions (**Figure S5 and S7**). gRNA sequences were given in **Supplementary Table 3**.

### Caspase inhibition, stress response inhibition, cholesterol biosynthesis inhibition, TGFβ signaling inhibition and mutant IDH1 inhibition assays

Cells were seeded on 96-well plates and treated with stated inhibitors the next day. For caspase inhibition assay, Z-FA-FMK (Negative Control) or Z-VAD-FMK (General Caspase Inhibitor) pre-treatments were performed at 20µM final concentration for 24h before additional drug treatments. For PERK and ISR inhibition, PERKi (1µM) or ISRIB (1µM) pre-treatments were performed for 24h before additional drug treatments. Cholesterol biosynthesis was inhibited using Simvastatin (50µM) or Lovastatin (50µM). They were given either individually or after 24h of ISRIB, or together with Belinostat. For inhibition of TGFβ signaling, RepSox (0.5µM) or SB-431542 (0.5µM), were given to cells either individually or together with Belinostat. For inhibition of mutant IDH1 enzyme, cells were pre-treated with GSK864 (2.5µM), a selective IDH1^R132H^ inhibitor^11^ for 3 days before any assay. Primary MGG119 and MGG152 cells were treated with GSK864 (2.5µM) for 10 passages to observe long term effects.

### Total cholesterol assay and rescue experiment with cholesterol or LDLR activation

Endogenous cholesterol levels in A172 and MGG152 cells were measured by Amplex Red Cholesterol Assay Kit (Thermo Fisher, USA) according to the manufacturer’s instructions. Details are described in **Supplementary Information**. For the rescue experiments, either parental cells, or cells with non-targeting or LDLR targeting gRNAs were seeded on 96-well plates as 4000 cells/well and treated with GSK-J4 in the presence or absence of exogenous cholesterol the next day (250µM, Sigma-Aldrich, USA). Cell viability was measured after 72h of the treatment.

### Live cell imaging

Olympus Xcellence Pro inverted microscope with 10X air objective (Center Valley, PA, USA) was used for the live-cell imaging experiments in a 37°C chamber supplied with 5% CO_2_. A172 wild type and IDH1^R132H^ cells were seeded on 24-well plates as 16.000 cells/well. Next day, they were treated with GSK-J4 (2.5µM) and/or Belinostat (1µM). Images of random positions from each well were captured with 8-10 min intervals during 96h after drug treatment. Cell numbers in 3 different frames for each condition was counted using ImageJ software (NIH Image, MD, USA) and viability curves were obtained for each condition.

### RNA sequencing

A172 wild-type and IDH1^R132H^-overexpressing cells were cultured in the presence or absence of GSK864 (2.5µM) for 3 days and cell pellets were collected for RNA isolation. MGG152 cells were cultured without drugs as control, with GSK-J4 or with Belinostat. Cell pellets were collected after 48h of drug treatments. All samples were studied as duplicate. Details of RNA isolation, library preparation and sequencing are described in **Supplementary Information**.

### qRT-PCR experiments

Cell pellets were collected after specified treatments and stored at - 80°C. Details are described in **Supplementary Information** and primers used in qRT-PCR experiments are listed in **Supplementary Table 2**.

### ChIP-qPCR experiments

Pellets were collected from MGG119 and MGG152 cells after 48h of GSK-J4 or Belinostat treatment. Details of Chromatin Immunoprecipitation are described in **Supplementary Information**.

### *In vivo* studies

Non-obese diabetic/severe combined immunodeficiency (NOD/SCID) mice were used for generation of orthotopic tumor models. All experiments were performed in Koç University Animal Facility with appropriate conditions and all protocols were approved by the Koç University Ethics Committee. IDH1 mutant primary MGG152 cells were infected with lentiviruses containing both Firefly Luciferase and mCherry. 2×10^5^ cells were injected intracranially using stereotaxic injection in 7µl PBS. To monitor tumor formation and progression, D-Luciferin (50mg/kg) was injected intraperitoneally, and luciferase activity was measured via in vivo bioluminescence imaging system (IVIS Lumina III). After 17 days of tumor injection, mice were treated with either DMSO or GSK-J4 and Belinostat combination. Both compounds were injected intraperitoneally as 100mg/kg daily, for 6 consecutive days. Tumor sizes were calculated as average radiance via Living Image software (PerkinElmer, USA). Kaplan-Meier survival plot was also generated via using GraphPad Prism 8.0.2, GraphPad Software, (CA, USA).

### Statistical analysis

All data were analyzed with GraphPad Prism or ImageJ software. Data were presented as mean +/-SD. For the comparison of two groups, unpaired t-test was used to determine the p-value. Additionally, one-way ANOVA was used for the comparisons including multiple parameters. A two-sided p-value<0.05 was considered statistically significant. Details of analyses were given in figure legends.

## RESULTS

### Epigenetic inhibitor screen identifies potent compounds targeting *IDH1*-mutant gliomas

We conducted a chemical screen in two independent primary glioma cell lines, MGG119 and MGG152, that carry an endogenous R132H point mutation in *IDH1*^21^, using a focused library of 46 compounds targeting different classes of chromatin modifiers (**Figure 1A**). DMSO-only or untreated cells served as negative controls; most compounds had minimal effect on cell viability (reducing viability to 92.4±15.3% for MGG119, and to 86.7±21.7% for MGG152) (**Figure 1B**). We considered a compound a “hit” if it reduced cell viability by 1 SD or lower (77.1% and 65.0%) in both cell lines. Accordingly, 5 compounds exhibited significant effects on both cell lines, namely 5-azacytidine; a DNA methyltransferase (DNMT) inhibitor, Chaetocin; an unspecific lysine-methyltransferase (KMT) inhibitor, GSK-J4; a histone H3K27me3 demethylase inhibitor, and Belinostat and Trichostatin A, histone deacetylase (HDAC) inhibitors. The effects of these compounds were dose-dependent on MGG119 and MGG152 cells; but were minimal on two non-malignant human fibroblast lines (Fibro1 and 2) (**Figure 1C**). To examine their combinatorial efficacy, we applied dual combinations of 4 screen hits targeting different types of chromatin modifiers in *IDH1*-mutant cells or fibroblasts (**Figure 1D**). Fibroblasts were not affected by compounds applied individually or in combination; however, *IDH1*-mutant glioma cells displayed significantly reduced viability upon combination treatments **(Figure S1A-E, Figure 1E**). Specifically, the combination of GSK-J4 and Belinostat showed synergistic effects against *IDH1*-mutant cell lines (**Figure 1F-G, Figure S1F-G)**. A GSK-J4 and Belinostat combination induced apoptosis in *IDH1*-mutant glioma cells, as demonstrated by markedly increased Poly (ADP-ribose) polymerase-1 (PARP) cleavage (**Figure 1H**).

**Figure 1.**
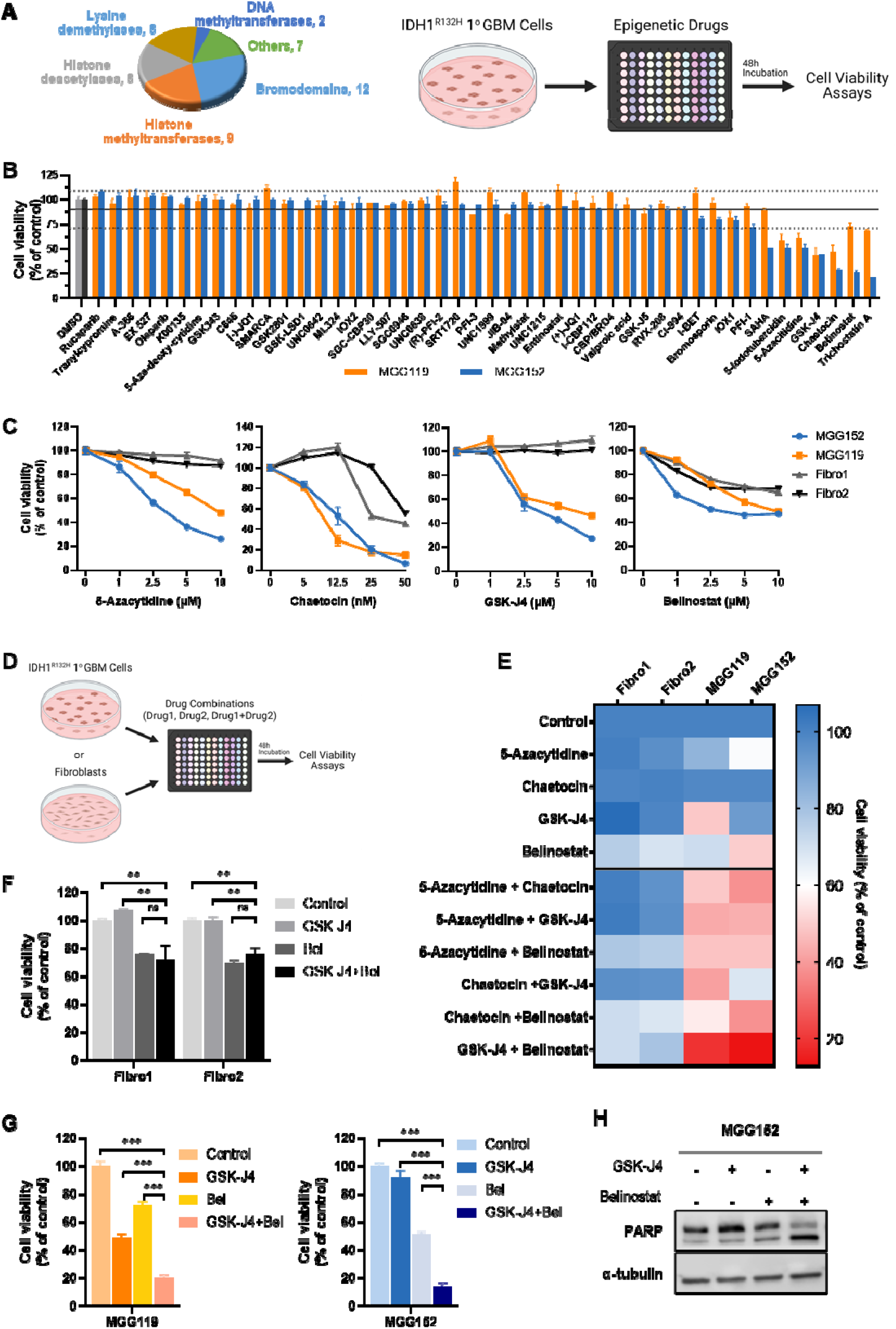
Epigenetic inhibitor screen identifies potential drugs for *IDH1*-mutant gliomas. **A)** Pie chart showing the total of 46 chemical probes targeting different classes of epigenetic enzymes and schematic representation of drug screen. **B)** Percent viability of patient-derived *IDH1*-mutant MGG119 and MGG152 cells treated with the chemical probe library for 48h. Black and gray bars represent DMSO control, the horizontal black line indicates the mean of viability, the staggered lines denote SD=1. **C)** Line plots showing the percent viability of patient-derived *IDH1*-mutant primary GBM cells or non-malignant fibroblasts treated with screen hits individually for 72h. **D)** Schematic view of the drug combination strategy for *IDH1*-mutant primary GBM cells and patient-derived fibroblasts. All possible dual combinations of screen hits were applied. **E)** Heatmap depicting the effects of individual and combinatorial treatments of screen hits (for 72h) on viability. Percent cell viability is expressed by a color scale (blue-red: viable-dead). **F, G)** Bar graphs indicating the viability of non-malignant fibroblasts **(F)** and *IDH1*-mutant MGG119 and MGG152 cells **(G)** upon treatment with GSK-J4 (2.5 µM) and/or Belinostat (1 µM) for 72h. **H)** Western blot images showing the alteration of PARP cleavage upon GSK-J4 and/or Belinostat treatment in *IDH1*-mutant MGG152 cells. α-tubulin is displayed as a loading control. P-values were determined by unpaired t-test; ns, non-significant; *p < 0.05; **p < 0.01; ***p < 0.001.

### GSK-J4 and Belinostat combination selectively targets *IDH1*-mutant glioma cells

To examine IDH1-mutation related effects of GSK-J4 or Belinostat further, we generated a paired cell line system via overexpression of mutant IDH1 (*IDH1*^*R132H*^) in A172 cells. Immunohistochemical staining and Western blot analysis confirmed the expression of mutant IDH1 (**Figure 2A-B**). The production of 2-HG was pronounced in the A172 *IDH1*^*R132H*^ cells, which could be reversed by 3-day treatment with GSK864, an inhibitor of mutant IDH1 enzyme **(Figure 2C)**. Testing the epigenetic inhibitors on this pair, we observed that A172 *IDH1*^*R132H*^ cells were significantly more sensitive to individual GSK-J4 or Belinostat treatments than A172 wild-type cells **(Figure 2D)**. This was also evident in another similarly established cell line pair **(Figure S2A-C)**. Together, these results demonstrate genotype-selective vulnerability of *IDH1*^*R132H*^ cells to GSK-J4 or Belinostat treatment. We next assessed the combinatorial efficacy of GSK-J4 and Belinostat and observed that a GSK-J4/Belinostat combination significantly reduced the viability of *IDH1*^*R132H*^ cells. In contrast, no synergistic effect of the compounds was observed in wild-type cells (**Figure 2E, S2D)**. We then utilized a live cell imaging system to examine the dynamics of GSK-J4 and Belinostat induced cell death (**Figure 2F-G, Supplementary videos 1-8)**. Quantification of the number of live cells at 0h, 24h, 48h, 72h and 96h demonstrated that *IDH1*^*R132H*^ cells were markedly more vulnerable to GSK-J4 and/or Belinostat treatment than wild-type cells (**Figure 2F**). While GSK-J4 or Belinostat mono-treatment halted growth of *IDH1*^*R132H*^ cells, their combination led to a significant induction of cell death (**Figure 2G)**. These potent selective effects of GSK-J4 and/or Belinostat treatment on *A172 IDH1*^*R132H*^ cells were further investigated with co-culture experiments of tumor cells and fibroblasts (**Figure S3A)**. In co-cultures of A172 *IDH1*^*R132H*^ and Fibro1 cells; fibroblasts remained viable but A172 *IDH1*^*R132H*^ cells were significantly reduced in number as measured by selective fluorescence labeling (**Figure S3B**). Similarly, normal human astrocytes (HA) were not affected by GSK-J4 and/or Belinostat treatment (**Figure S3C**). Interestingly, pretreatment of A172 *IDH1*^*R132H*^ cells with GSK864 reversed the selective sensitivity to the GSK-J4 and Belinostat combination (**Figure 2H)**. Similarly, drug sensitivity of MGG152 cells could be partly abrogated after long-term passaging with GSK864 (**Figure S2E**). Together, these results indicate that the combined treatment of GSK-J4 and Belinostat is selective and effective against *IDH1*-mutant glioma cells.

**Figure 2.**
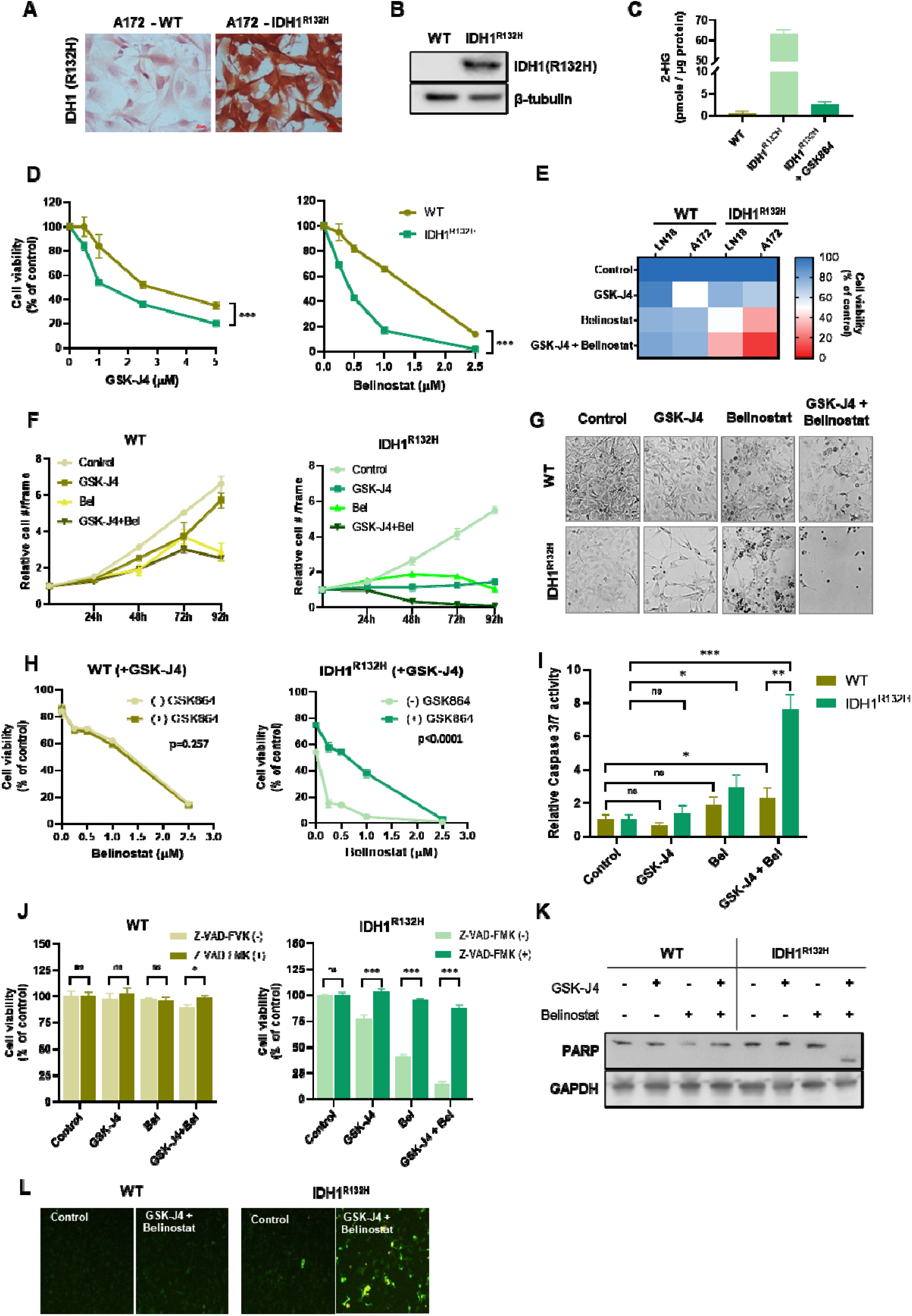
GSK-J4 and Belinostat combination selectively targets IDH1^R132H^ cells *in vitro*. Mutant IDH1 enzyme (*IDH1*^*R132H*^) was overexpressed in A172 GBM cells to generate wild-type (WT) and mutant (*IDH1*^*R132H*^) cell pairs. **(A)** Immunohistochemical staining of A172 cells that are WT or overexpressing IDH1^R132H^. Scale bars=20 µm. **B)** Western blot image showing the expression level of mutant IDH1 enzyme in A172 cell pair. β-tubulin is displayed as a loading control. **C)** Bar plot indicating the level of 2-HG in both A172 WT and *IDH1*^*R132H*^, and in *IDH1*^*R132H*^ cells treated with GSK864 (2.5 µM), an inhibitor of the mutant IDH1 enzyme. **D)** Line graphs depicting the viability of WT and *IDH1*^*R132H*^ cells after 72h treatment with the indicated doses of GSKJ4 or Belinostat. **E)** Heatmap displaying the effects of GSK-J4 and Belinostat, individually or in combination, on the viability of WT and *IDH1*^*R132H*^ pairs of A172 and LN18 GBM cells. Percent cell viability is expressed by a color scale (blue-red: viable-dead). **F)** Quantification of live cell images of A172 WT and *IDH1*^*R132H*^ cells after treatment of control, GSK-J4 (2.5 µM), Belinostat (1 µM), or their combination at 24h, 48h, 72h, and 92h. **G)** Representative live-cell images of A172 WT and *IDH1*^*R132H*^ cells after 92h treatment. **H)** Line plots showing the percent viability of GSK864 (2.5 µM) pre-treated or untreated A172 WT and *IDH1*^*R132H*^ cells upon treatment with the four different concentrations of Belinostat in the presence of GSK-J4 (2.5 µM). **I)** Bar plot indicating relative Caspase 3/7 activity in A172 WT and *IDH1*^*R132H*^ cells after 48h individual and combination treatments of GSK-J4 and Belinostat. **J)** Graphs demonstrating the effect of Z-VAD-FMK pre-treatment (20 µM for 24h), a general caspase inhibitor, on the viability of the A172 cell pair treated with GSK-J4 and/or Belinostat. **K)** Western blot showing the alterations of PARP cleavage in A172 cell pair upon GSK-J4 and/or Belinostat treatments. GAPDH is displayed as a loading control. **L)** Representative YO-PRO1 images showing cells undergoing apoptosis in the A172 WT and *IDH1*^*R132H*^ cells upon GSK-J4 + Belinostat treatment for 48h. (Green indicates cells undergoing apoptosis). For panel D and H, p-values were determined by 2-way ANOVA whereas for all other panels, p-values were determined by unpaired t-test; ns, non-significant; *p < 0.05; **p < 0.01; ***p < 0.001.

We then investigated apoptosis markers in wild-type and *IDH1*^*R132H*^ cells. While GSK-J4 or Belinostat as single agents slightly increased Casp3/7 activity, combinatorial treatment resulted in significant elevation of Casp3/7 activity in A172 *IDH1*^*R132H*^ cells (**Figure 2I**). Application of the general caspase inhibitor Z-VAD-FMK prevented GSK-J4 and Belinostat induced cell death (**Figure 2J)**. Cleavage of PARP-1 and apoptotic cell number was enhanced upon combinatorial treatment in A172 *IDH1*^*R132H*^ cells, but not in wild-type cells (**Figure 2K-L**). Together, these results suggest that *IDH1*-mutant glioma cells are more vulnerable to GSK-J4 and Belinostat treatment, and this vulnerability is dependent on mutant IDH1 activity, as its inhibitor reversed this phenotype.

### IDH1^R132H^ overexpression leads to global transcriptomic alterations, decreased cholesterol synthesis and increased stress response

To define molecular differences between the wild-type and *IDH1*^*R132H*^ cells, cells were grown with or without GSK864 and their transcriptomic differences were examined by RNA sequencing. **(Figure 3A)**. *IDH1*^*R132H*^ cells displayed a high number of differentially expressed genes (DEGs) when compared to wild-type cells. Specifically, there were 463 DEGs between wild-type and *IDH1*^*R132H*^ cells; and 86 DEGs in the presence of GSK864 (|log2|>1, p<0.05) **(Figure S4A-B)**. Wild-type cells were not affected by GSK864 as there were only 2 DEGs with GSK864 treatment. (**Figure S4A-B**). Heat maps and cluster diagrams revealed significant differential expression between the wild-type and *IDH1*^*R132H*^ cells, which was reversible upon GSK864 treatment (**Figure 3A**). Gene Set Enrichment Analysis (GSEA)^22^ demonstrated that several important pathways were activated in *IDH1*^*R132H*^ cells compared to wild-type cells (**Figure 3B-C)**. These pathways included several inflammation related networks, such as “Interferon alpha response”, “TNFα signaling via NFKB”, “Interferon gamma response”, “Inflammatory response”, and cell stress-related networks such as “Response of eIF2AK4/GCN2 to amino acid deficiency”, “Response of eIF2AK1/HRI to heme deficiency”, “PERK regulates gene expression” and “ATF4 activates genes in response to endoplasmic reticulum stress”. Enrichment of these pathways was reversed in the presence of GSK864 in *IDH1*^*R132H*^ cells (**Figure 3B-C**). Similarly, pathways that were inhibited in *IDH1*^*R132H*^ cells compared to wild-type cells, especially those related to lipid and cholesterol biosynthesis were reactivated in the presence of GSK864 (**Figure 3B-C**). A heat map showing the log2fold change of genes involved in cholesterol biosynthesis, interferon alpha response, and ATF4-mediated stress response was consistent with GSEA results (**Figure 3D**). qRT-PCR analysis confirmed the upregulation or downregulation of selected genes involved in these pathways in *IDH1*^*R132H*^ cells, which were reversible with GSK864 treatment (**Figure 3E-F, S4C-D**). In parallel with reduced gene expression, total cholesterol levels were markedly decreased in A172 *IDH1*^*R132H*^ cells compared to wild-type cells (**Figure 3G**). Moreover, *IDH1*^*R132H*^ cells were more sensitive to Thapsigargin, an endoplasmic reticulum stress inducer, and this phenotype could be reversed by GSK864 (**Figure 3H**). Thus, the wild-type and *IDH1*^*R132H*^ A172 cell line pair enabled studying the effects of *IDH1* mutation in glioma and revealed *IDH1* mutation induced global transcriptome changes that highlighted decreased cholesterol synthesis and increased stress response in a 2-HG dependent manner (**Figure 3I**).

**Figure 3.**
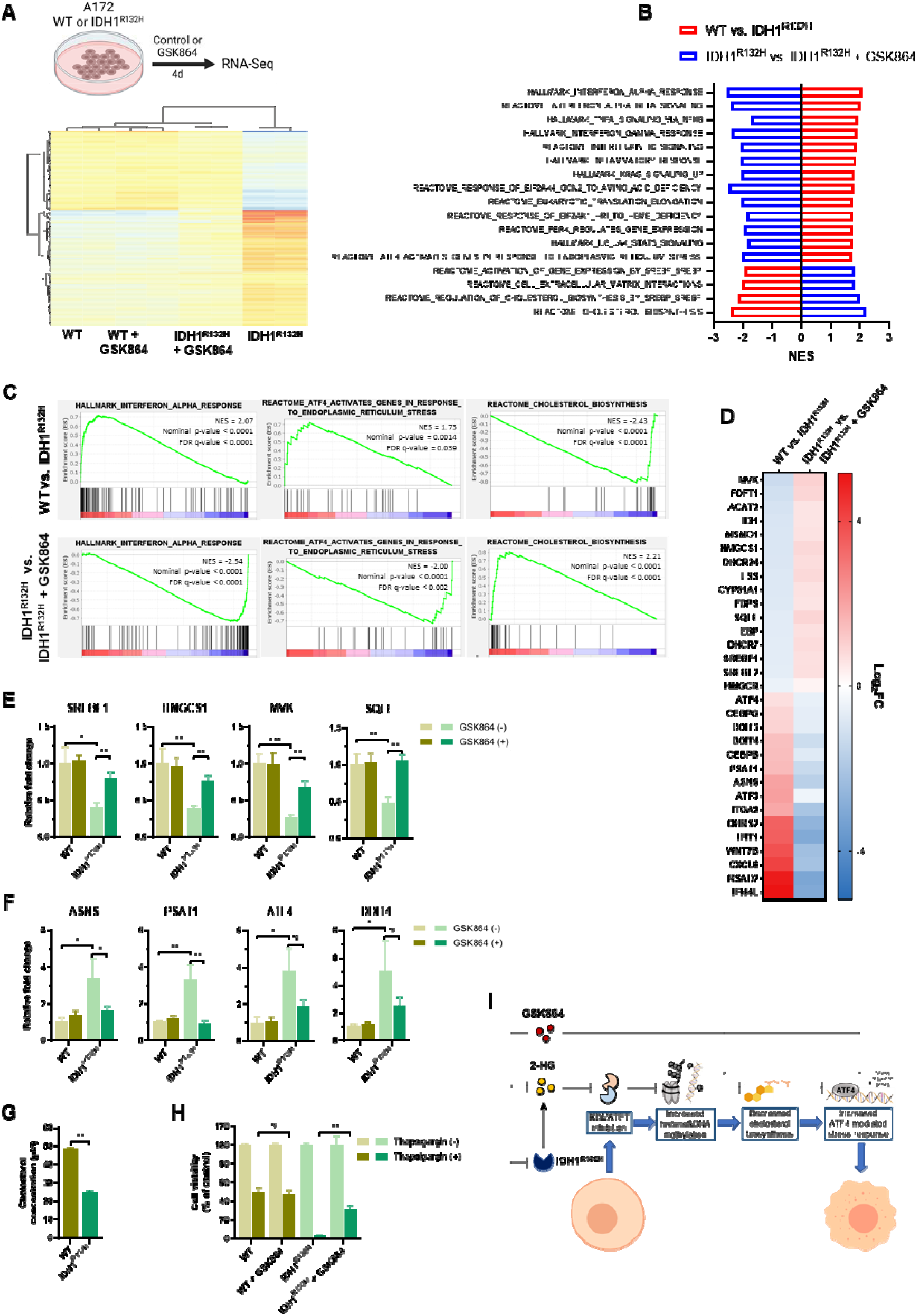
IDH1^R132H^ overexpression leads to global transcriptome alterations, decreased cholesterol synthesis and increased stress response that are reversible with GSK864. **A)** Schematic representation of GSK864, an inhibitor of mutant IDH1, treatment on A172 WT and *IDH1*^*R132H*^ cells for RNA-Seq analysis. Heatmap of differentially expressed genes (DEGs) between WT and *IDH1*^*R132H*^ cells with or without GSK864. **B)** Bar graph showing the top enriched, both negatively and positively, Reactome and Hallmark pathways upon *IDH1*^*R132H*^ overexpression and GSK864 treatment on *IDH1*^*R132H*^ cells. **C)** GSEA enrichment plots demonstrating relative expression changes in genes involved in the top indicated cellular processes (B) in A172 WT vs *IDH1*^*R132H*^ cells and in *IDH1*^*R132H*^ cells with vs without GSK864 treatment. **D)** Heatmap displaying Log2 fold change of genes involved in the enriched pathways (C). **E, F)** Validations of selected RNA-Seq results via qRT-PCR. **G)** Bar graph showing cellular cholesterol concentration in A172 WT and *IDH1*^*R132H*^ cells. **H)** Bar graph indicating the effects of Thapsigargin, an ER stress inducer, on A172 cell viability either in the presence or absence of GSK864. NES, normalized enrichment score. FDR, false discovery rate. For panel E, F, G, and H the p-values were determined by unpaired t-test; ns, non-significant; *p < 0.05; **p < 0.01; ***p < 0.001. **I)** Model describing the differences between WT and *IDH1*^*R132*^ cells.

### GSK-J4 inhibits cholesterol biosynthesis and activates ATF4-mediated stress response in *IDH1*-mutant cells

GSK-J4 is an inhibitor of KDM6A/UTX and KDM6B/JMJD3, which are H3K27 demethylases^23^, with some activity towards KDM5B and KDM5C; therefore, we investigated its specificity. GSK-J5, an inactive form of GSK-J4, had no effect on *IDH1*-mutant cells (**Figure S5A-B**). KDM5-specific inhibitors^24^, KDM5-C70 and KDOAM25a, or pan-2-OG inhibitor IOX-1, were also ineffective on *IDH1*-mutant cells (**Figure S5C-D**). In contrast, KDOBA67, another compound targeting KDM6 enzymes^25^, displayed similar efficacy compared to GSK-J4 (**Figure S5C-D**). Targeting *KDM6A* and/or *KDM6B* genes via CRISPR/Cas9 demonstrated that ablation of both KDM6A and KDM6B phenocopied the effects observed with GSK-J4 in *IDH1*-mutant cells (**Figure S5E-G**).

GSK-J4 treatment slightly increased the global H3K27me3 levels in MGG152 cells (**Figure 4A**). To examine the effects of GSK-J4 on transcription of MGG152 cells, we performed RNA sequencing (**Figure 4B**). GSEA analysis indicated many transcriptional networks were deregulated upon GSK-J4 treatment (**Figure 4C**). Namely, lipid and cholesterol biosynthesis pathways were downregulated, and integrated stress response (ISR) related pathways, such as “ATF4-mediated stress response” were upregulated (**Figure 4D**). Volcano plots indicated the differential expression of genes involved in these pathways (**Figure 4E**), which was further validated by qRT-PCR in both MGG152 and MGG119 cells (**Figure 4F, S6A**). To investigate whether these transcriptomic alterations were caused by direct effects of GSK-J4 on histone methylation, we performed ChIP-qPCR. Accordingly, GSK-J4 increased H3K27me3 levels in the promoter of genes involved in cholesterol biosynthesis (**Figure 4G, S6B**). Already low total cholesterol levels in *IDH1*-mutant cells were further decreased upon GSK-J4 treatment, which is recovered by CRISPR-mediated LDLR activation (**Figure S6D, 4H**). Exogenous cholesterol treatment (**Figure S6C**) or LDLR activation (**Figure 4I**) rescued GSK-J4-induced decrease in cell viability in *IDH1*-mutant cells.

**Figure 4.**
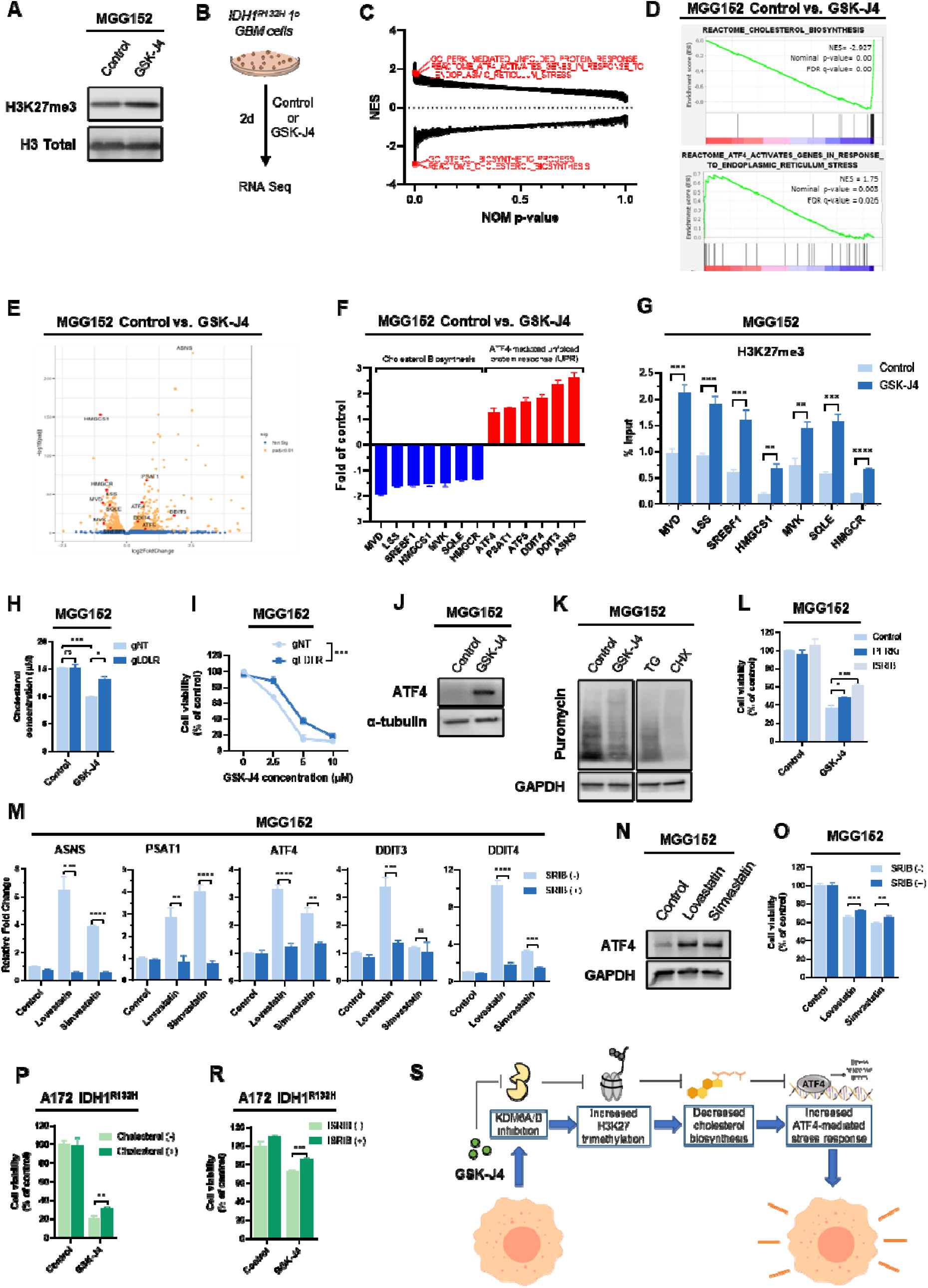
GSK-J4 inhibits cholesterol biosynthesis and activates ATF4-mediated stress response in IDH1-mutant GBM cells. **A)** Western blot images showing global change in H3K27 tri-methylation upon 48h of GSK-J4 treatment (2.5 µM). **B)** Schematic representation of sample preparation for RNA-seq. **C)** GSEA of MGG152 cells treated with GSK-J4 using pre-ranked 780 gene lists according to log2FC value for comparisons, for all gene sets available at The Molecular 782 Signatures Database (MSigDB). **D)** GSEA plots indicating upregulated and downregulated pathways upon GSK-J4 treatment. NES=normalized enrichment score, FDR=false discovery rate. **E)** Volcano plot showing differentially expressed genes (DEGs) upon GSK-J4 treatment. Genes having role in cholesterol synthesis and ATF4-mediated stress response pathways are indicated with red dots. **F)** Validation of RNA-seq results for genes involved in cholesterol synthesis and ATF4-mediated stress response pathways upon 48h of GSK-J4 treatment. **G)** ChIP-qPCR analysis of relative H3K27me3 occupancy in the promoter regions of genes involved in cholesterol biosynthesis pathway, with and without GSK-J4 treatment. **H)** Total cholesterol concentration in MGG152 cells infected with non-targeting (gNT) or LDLR targeting (gLDLR) gRNAs together with dCas9-TET1 and MPH viruses, with or without GSK-J4 treatment. **I)** Relative viability of either control (gNT) or LDLR actiated (gLDLR) MGG152 cells with different doses of GSK-J4 treatment for 48h. **J)** Western blot image showing ATF4 protein expression upon 48h of GSK-J4 treatment. **K)** SUnSET assay showing global translation levels after 48h of GSK-J4 treatment in MGG152 cells. Thapsigargin (TG, 5 µM) or cycloheximide (CHX, 5 µM) were used as positive controls. **L)** Relative cell viability after 48h of GSK-J4 treatment in the presence or absence of PERKi (1 µM) or ISRIB (1 µM). **M)** Relative expression of genes involved in ATF4-mediated stress pathway upon lovastatin (50 µM) or simvastatin (50 µM) treatment in the presence or absence of ISRIB (1 µM). **N)** Western blot image showing effect of lovastatin or simvastatin treatment on ATF4 protein levels. **O)** Relative viability of MGG152 cells after 48h of lovastatin or simvastatin treatment in the presence or absence of ISRIB (1 µM). **P)** Relative viability of A172 IDH^R132H^ cells after 48h of GSK-J4 treatment in the presence or absence of exogenous cholesterol (250 µM). **R)** Relative viability of A172 IDH^R132H^ cells after 48h of GSK-J4 treatment in the presence or absence of ISRIB (1 µM). **S)** Schematic model of intracellular changes upon GSK-J4 treatments in IDH1-mutant GBM cells. For panel I, p-value was determined by 2-way ANOVA whereas for all other panels, p-values were determined by unpaired t-test; ns, non-significant; *p < 0.05; **p < 0.01; ***p < 0.001, ****p<0.0001.

Given that ATF4 is a major transcription factor that regulates survival and apoptosis under various stress conditions^26^, we examined the effects of GSK-J4 on ATF4 activation. GSK-J4 treatment increased ATF4 protein levels (**Figure 4J**) and decreased global translation rate, which is a cell stress indicator^27^, (**Figure 4K**). Inhibition of ISR either by PERK inhibitor (PERKi) or ISRIB, significantly recovered the cell viability (**Figure 4L**). Linking cholesterol de-regulation and stress response, chemical inhibitors of cholesterol biosynthesis, lovastatin or simvastatin, induced ATF4-mediated stress response genes, which were potently blocked by ISRIB (**Figure 4M, S6E**). Lovastatin or simvastatin markedly increased ATF4 protein levels (**Figure 4N, S6F**), and reduced cell viability, which was partially recovered by ISRIB in both MGG152 and MGG119 cells (**Figure 4O, S6G**). In parallel with the link between these pathways, exogeneous cholesterol or ISRIB partially blocked the effects of GSK-J4 on A172 *IDH1*^*R132H*^ cell viability (**Figure 4P-R**). Together, these results suggest that GSK-J4 inhibits the cholesterol biosynthesis pathway, which is linked with the activation of ATF4-mediated ISR in *IDH1*-mutant cells **(Figure 4S)**.

### Belinostat activates TGF-β and cholesterol efflux pathways and induces a G2/M cell cycle arrest in *IDH1*-mutant cells

Consistent with its pharmacological activity, Belinostat treatment increased H3K27 and H4 acetylation (**Figure 5A**). GSEA analysis indicated that Belinostat deregulated many pathways in MGG152 cells (**Figure 5B-C**). TGF-β signaling and cholesterol efflux pathways were among the top activated pathways, while pathways regulating cell cycle were inhibited upon Belinostat treatment (**Figure 5D**). Genes involved in these pathways were shown to be upregulated or downregulated in a volcano plot (**Figure 5E**). qRT-PCR showed that *ATF5*, which promotes survival under stress conditions^28^, anti-apoptotic *BIRC5*/survivin and *BCL2L1*/Bcl-xL were downregulated. Conversely, key genes involved in TGF-β signaling (*TGFB1* and *TGFBR2*), cholesterol efflux (*APOE* and *NPC2*), and cell cycle arrest (*CDKN1A*/p21) were upregulated (**Figure 5F, S6H**). ChIP-qPCR analyses revealed that H3K27 acetylation was deposited at promoters of these upregulated genes in both MGG152 and MGG112 cells (**Figure 5G, S6I**). In accordance with the activation of TGF-β signaling upon Belinostat treatment, inhibitors of TGF-β signaling, RepSox and SB-431542, recovered Belinostat-induced reduction of cell viability (**Figure 5H, S6J**). p21 protein levels were markedly increased upon Belinostat treatment, alongside an increase in acetylated α-tubulin (**Figure 5I**). Consistent with GSEA results and p21 induction, Belinostat induced a G2/M arrest in MGG152, MGG119 and A172 *IDH1*^*R132H*^ cells (**Figure 5J, S6K**). Further, statin and Belinostat combinations showed significant cooperation and potently reduced the viability of *IDH1*-mutant cells (**Figure 5K, S6L**). Together, these results suggest that HDAC inhibition by Belinostat activates TGF−β signaling and causes p21-mediated cell cycle arrest in *IDH1*-mutant cells (**Figure 5L)**.

**Figure 5.**
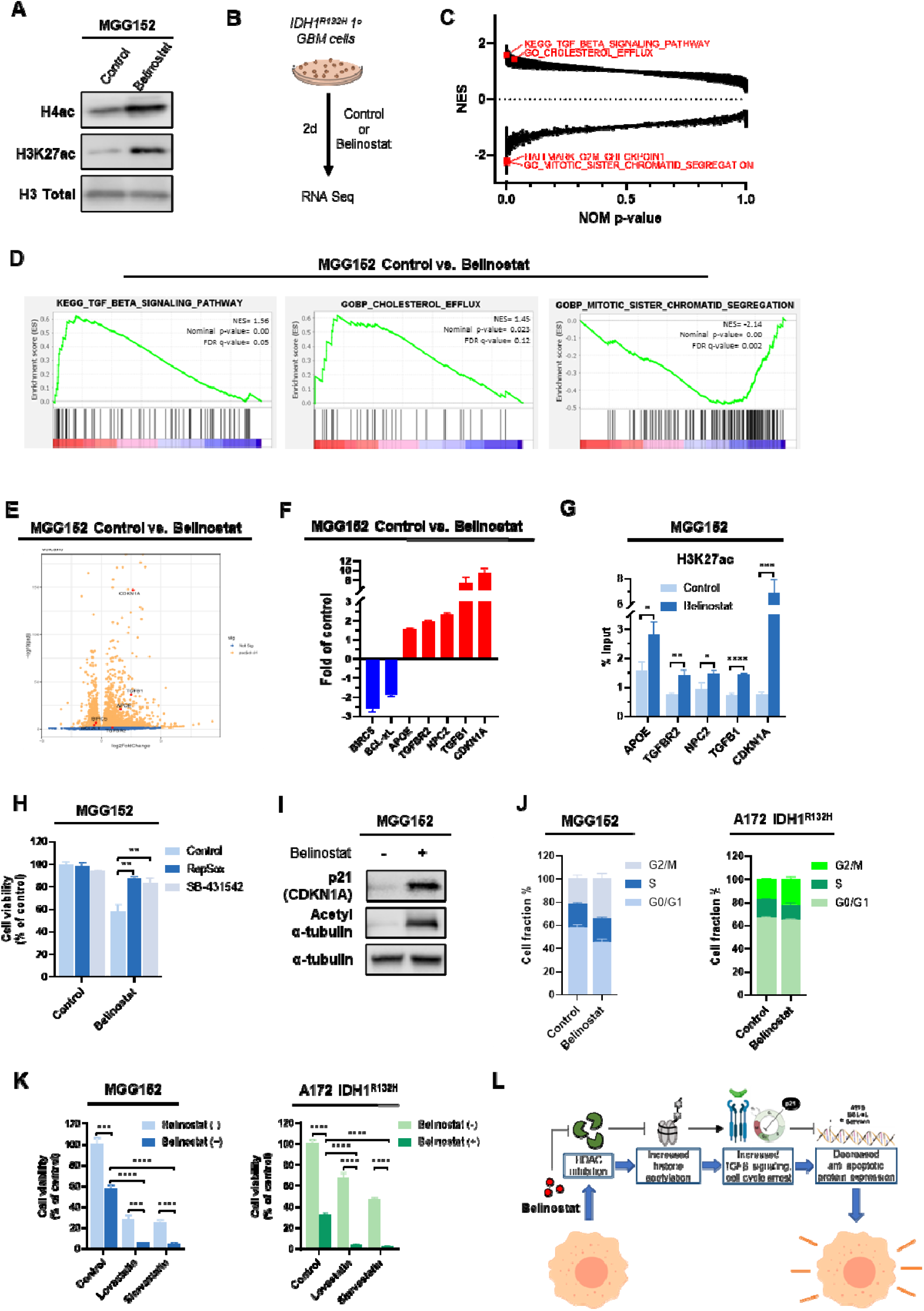
Belinostat activates p21-mediated cell cycle arrest and TGF-β pathway in IDH1-mutant GBM cells. **A)** Western blot images showing global changes in histone acetylation upon 48h of Belinostat treatment in MGG152 cells. Total H3 protein levels were used as loading control. **B)** Schematic representation of sample preparation for RNA-seq. **C)** GSEA of MGG152 cells treated with Belinostat using pre-ranked 780 gene lists according to log2FC value for comparisons, for all gene sets available at The Molecular 782 Signatures Database (MSigDB). **D)** GSEA plots derived from RNA-seq analysis in MGG152 cells indicating upregulated and downregulated pathways upon 48h of Belinostat treatment. NES=normalized enrichment score, FDR=false discovery rate. **E)** Volcano plot showing differentially regulated genes (DEGs) upon Belinostat treatments. Genes involved in TGF-β, cholesterol efflux and apoptosis/cell cycle pathways are indicated with red dots. **F)** Validation of RNA-seq results for genes involved in TGF-β, cholesterol efflux and apoptosis/cell cycle pathways. **G)** Relative H3K27ac occupancy in the promoter regions of genes involved in TGF-β, cholesterol efflux and apoptosis/cell cycle pathways, as measured by ChIP-qPCR. **H)** Relative viability of MGG152 cells after 48h of Belinostat treatment in the presence or absence of TGF-β inhibitors, RepSox (0.5 µM) or SB-431542 (0.5 µM). **I)** Western blot images showing changes in the protein level of p21 and acetyl-α-tubulin upon 48h of Belinostat treatment in MGG152 cells. α-tubulin protein levels were used as loading control. **J)** Graphs representing changes in cell cycle phases after 24h of Belinostat treatment in MGG152 and A172 IDH1^R132H^ cells. **K)** Relative viability of MGG152 and A172 IDH1^R132H^ cells after 48h of Lovastatin and Simvastatin treatments in the presence or absence of Belinostat (1 µM). **L)** Schematic model of intracellular changes upon Belinostat treatment in IDH1-mutant GBM cells. For all panels, p-values were determined by unpaired t-test; ns, non-significant; *p < 0.05; **p < 0.01; ***p < 0.001, ****p<0.0001.

### GSK-J4 and Belinostat combination induces apoptosis and inhibits *in vivo* growth of *IDH1*-mutant glioma

Severe and long-lasting stress can induce cell death^29^, therefore we investigated apoptosis pathway upon GSK-J4 and Belinostat combination treatment. Expression of *DDIT3*/CHOP, a pro-apoptotic stress response gene, and of *BBC3*/PUMA and *PMAIP1*/NOXA, transcriptional targets of CHOP^29^, were increased upon treatment (**Figure 6A-B**); and GSK864 reversed these changes (**Figure 6B**). To validate the role of *DDIT3*/CHOP in GSK-J4 and Belinostat-induced apoptosis, we knocked out *DDIT3* gene via CRISPR/Cas9 (**Figure S7**). *DDIT3*-knockout did not affect viability, however it significantly protected *IDH1*^*R132H*^ cells from GSK-J4 and Belinostat (**Figure 6C**). Induction of PUMA and NOXA by GSK-J4 and Belinostat treatment was potently inhibited in *DDIT3-*knockout cells (**Figure 6D**). To follow up on the decreased levels of Bcl-xL upon Belinostat alone or together with GSK-J4 (**Figure 5F and 6E**), we demonstrated that ectopic overexpression of Bcl-xL rescued the decreased viability of cells upon GSK-J4 and Belinostat treatment in *IDH1*-mutant cells (**Figure 6F**). Together, these results suggest that GSK-J4 and Belinostat-induced cell death involves the activation of stress-induced pro-apoptotic programs and inhibition of anti-apoptotic programs in *IDH1*-mutant cells.

**Figure 6.**
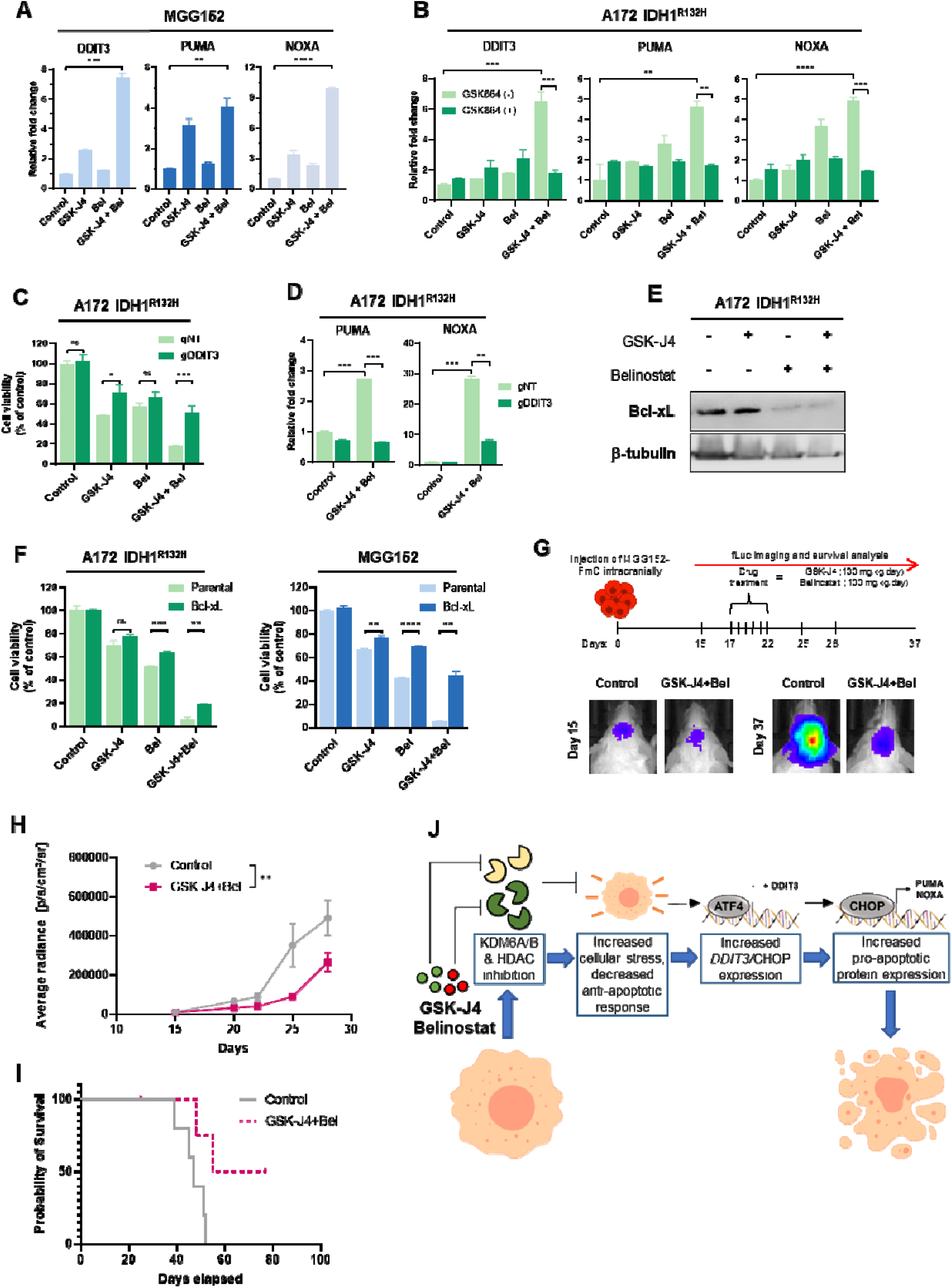
GSK-J4 and Belinostat combination induces DDIT3-mediated apoptosis and reveals efficacy in IDH1-mutant GBM cells *in vivo*. **A)** Relative expression of *DDIT3, PUMA* and *NOXA* genes in MGG152 cells upon treatment with GSK-J4, Belinostat or their combination. **B)** Relative expression of *DDIT3, PUMA* and *NOXA* genes in A172 IDH1^R132H^ cells upon treatment with GSK-J4, Belinostat or their combination in the presence or absence of mutant IDH1 inhibitor, GSK864. **C)** Viability of A172 IDH1^R132H^ cells expressing non-targeting (gNT) or DDIT3 (gDDIT3) gRNA upon treatment with GSK-J4 and/or Belinostat. **D)** Relative expression of PUMA and NOXA genes in A172 IDH1^R132H^ cells expressing non-targeting (gNT) or DDIT3 (gDDIT3) gRNA upon treatment with GSK-J4 and Belinostat combination. **E)** Western blot image showing Bcl-xL protein levels upon treatment with GSK-J4, Belinostat or their combination. β-tubulin was used as loading control. **F)** Viability of parental or Bcl-xL overexpressing A172 IDH1^R132H^ or MGG152 cells upon treatment with GSK-J4, Belinostat or their combination. For panels A-F, p-values were determined by unpaired t-test; ns, non-significant; *p < 0.05; **p < 0.01; ***p < 0.001, ****p<0.0001 **G)** Schematic representation for *in vivo* experiments conducted with MGG152 cells, expressing Fluc and mCherry (FmC) (top). Representative bioluminescence images of intracranial tumors on days 15 and 37 (bottom). **G)** Average radiances [p/s/cm^²^/sr] of control or drug combination treated tumors. p-value was calculated via two-way ANOVA test, *p < 0.05; **p < 0.01; ***p < 0.001. **H)** Kaplan-Meier survival plot for control or drug treated mice (n=5). p-value was calculated via Log-rank test (p=0.0318). **I)** Schematic model of intracellular changes upon GSK-J4 and Belinostat combination treatment in IDH1-mutant GBM cells.

We next tested the therapeutic potential of a GSK-J4 and Belinostat combination in an intracranial xenograft model derived from Luciferase-expressing MGG152 cells. When the tumors reached detectable volumes, we administered GSK-J4 and Belinostat or DMSO control solutions intraperitoneally for 6 days (**Figure 6G**). GSK-J4 and Belinostat co-treatment significantly decreased tumor volume compared to the control group (**Figure 6H)**. Prior to drug treatment, on day 15, average tumor volumes were comparable between the animals, but at the end of the treatment period, the average tumor volume was significantly lower in the drug-treated group. This difference was maintained until day 28 (**Figure 6H**). Kaplan-Meier survival plots indicated significantly longer survival of animals treated with the combination regimen, with half of the mice surviving long-term (**Figure 6I**). Taken together, these results demonstrate the combined efficacy of GSK-J4 and Belinostat on orthotopic growth of *IDH*-mutant glioma *in vivo*.

## DISCUSSION

We interrogated the epigenetic vulnerabilities of *IDH*-mutant gliomas with a chemical screen and identified a potent epigenetic drug combination to treat *IDH1*-mutant glioma cells. The GSK-J4 and Belinostat combination exhibited a powerful synergistic effect on *IDH1*-mutant cells with minimal effects on *IDH1*-wild-type or non-malignant cells. In the screen with inhibitors of several chromatin modifiers, such as HDACs, HATs, HMTs, KDMs, PHDs, DNMTs, methyl lysine binders, and Bromodomain proteins, we identified 5 compounds (5-azacytidine, Chaetocin, GSK-J4, Belinostat and TrichostatinA) with anti-proliferative effects in primary *IDH1*-mutant cells. Among them, 5-azacytidine was previously shown to be effective against *IDH*-mutant glioma^30^. Indeed, there are ongoing clinical trials (NCT02677922, NCT03684811) for 5-azacytidine treatment alone or combined with mutant IDH inhibitors. Recently, SAHA, another HDAC inhibitor, was shown to be effective on mutant IDH1 overexpressing glioma cell lines, regulating DNA repair pathways^31^. To our knowledge, our study is the first to identify new epigenetic drugs targeting *IDH1*-mutant cells in a screen platform using patient-derived, endogenously *IDH1*-mutant glioma cells.

*IDH1* mutation is an early genetic event in LGG tumorigenesis and is conserved in recurrent tumors^32^. Inhibitors of mutant IDH1 enzyme have been prime clinical candidates^10–12^ and we here show that mutant IDH1 related gene expression programs constitute a selective vulnerability for GSK-J4 and Belinostat, which is reversible with mutant IDH1 inhibitor, GSK864. The metabolic reprogramming through IDH1 mutation is profound, where induction of cell stress and apoptosis related pathways were partly reversed by GSK864 and are consistent with previous reports on the effects of mutant IDH inhibitors^13–15^. Collectively, our work suggests that in addition to directly blocking mutant IDH1 activity in tumors, the exploitation of selective vulnerabilities caused by the *IDH1*-mutation might constitute an additional avenue for future clinical translation.

Our transcriptomic analysis of paired cell lines reveal reduced cholesterol biosynthesis in *IDH1*-mutant cells, consistent with previous observations of low cholesterol and lipid in *IDH1*-mutant cells^33^. We also observed ATF4-mediated ISR activation in *IDH1*-mutant cells, which may be an adaptive response to decreased lipid and cholesterol levels, in accordance with previous work linking cholesterol biosynthesis dysregulation and ISR^34^. However, there may be other mechanisms responsible for the ISR phenotype in *IDH1*-mutant cells. A recent report showed that 2-HG inhibits prolyl hydroxylase and causes ER stress via immature collagen accumulation in *Idh1-*R132H knock-in mice^35^. Another study demonstrated that 2-HG alters glutamine metabolism and increases mitochondrial stress^7^. Similarly, immune response pathways and genome instability were shown to be upregulated in *IDH*-mutant cells, as a result of activation of endogenous retroviruses (ERVs)^36^. ERV activation was also shown to induce ER stress and UPR^37^. Our transcriptomic analyses of paired wild-type and *IDH1*^*R132H*^ cells revealed similar results, where IFN-α, IFN-γ, inflammatory response pathways and ATF4-mediated ISR were activated in *IDH1*^*R132H*^ cells, which were all reversed with GSK864. Therefore, we suggest that *IDH1*-mutant cells have lower *de novo* cholesterol biosynthesis and higher basal cellular stress, up to levels that the cells can tolerate. However, GSK-J4 and Belinostat treatment can increase cell stress above this tolerable threshold, leading to subsequent cell death.

We conclude that GSK-J4, an inhibitor of KDM6A/B, exacerbates the metabolic vulnerabilities caused by *IDH1* mutation. The increase in global H3K27 methylation levels upon GSK-J4 treatment was small, possibly due to the basal hypermethylated state in *IDH1*-mutant cells. However, GSK-J4 might display loci-specific actions as we observed increased H3K27me3 levels at the promoters of genes involved in lipid and cholesterol biosynthesis. This is consistent with a study that offered KDM6A targeting as a potential therapeutic strategy for prostate cancer, through inhibition of lipid metabolism^38^. In accordance with the reported connection between dysregulated cholesterol synthesis and cellular stress^34^, we observed an ISR activation with ATF4-mediated changes upon GSK-J4 treatment in *IDH1*-mutant cells. GSK-J4-mediated ISR activation was also documented recently in various cancer cell lines^39,40^. To further establish a link between cholesterol biosynthesis and ISR, we blocked cholesterol synthesis using statins and observed activation of ATF4 and ISR. Conversely, exogenously administered cholesterol or increased cholesterol intake by LDLR upregulation provided a partial recovery. Therefore, suppression of cholesterol synthesis pathway genes through H3K27 hypermethylation and induction of ATF4-mediated ISR underlie the anti-tumor effects of GSK-J4 in *IDH1*-mutant cells.

In parallel, we showed that Belinostat increased global acetylation of histone H3 and H4 in *IDH1*-mutant cells. H3K27ac levels of the *CDKN1A*/p21 promoter were increased with Belinostat, in line with previous studies reporting p21 as an HDAC target^41^. Together with the dramatic p21 induction, we observed a G2/M cell cycle arrest in *IDH1*-mutant glioma cells, similar to previous reports in different cancers^42^. Besides p21 activation, expression of genes involved in TGF-β and cholesterol efflux pathways was induced along with higher H3K27ac occupancy at their promoters. Concomitantly, anti-apoptotic *ATF5, BIRC5*/survivin and *BCL2L1*/Bcl-xL expression was downregulated, possibly through indirect regulation. TGFβ signaling can have tumor suppressive effects^43^, and induction of TGFβ signaling by various HDAC inhibitors was previously observed and deemed responsible for repression of *BIRC5*/survivin^44^ and *BCL2L1*/Bcl-xL^45^. Indeed, TGFβ inhibitors significantly inhibited Belinostat-induced cell death in our *IDH1*-mutant cells. Interestingly, TGFβ signaling was previously shown to induce cholesterol efflux and reduce cholesterol intake^46^. Therefore, the use of statins together with Belinostat led to a significant decrease of *IDH1*-mutant cell viability, comparable to the GSK-J4 and Belinostat combination.

The synergistic effects of GSK-J4 and Belinostat treatment on *IDH1*-mutant cells could occur through multiple mechanisms. Future work is clearly needed to understand in detail the global epigenomic consequences of IDH1 mutations and the effects of HDAC and KDM6 inhibition. However, our work shows that transcriptional changes accompany an ISR in *IDH1*-mutant cells. ATF4 upregulated survival pathways under stress conditions, or apoptotic pathways under severe stress^26^ and GSK-J4 induced ATF4-mediated ISR and further reduced viability as cellular stress was already high in *IDH1*-mutant cells. However, with low doses of GSK-J4, expression of *ATF5*, a target of ATF4 responsible for survival pathways under stress conditions^28^, was induced as an adaptive mechanism. By combining with Belinostat, anti-apoptotic pathways were suppressed either by downregulation of Bcl-xL and survivin, or p21-mediated cell cycle arrest, driving the cells towards apoptosis. As Bcl-xL downregulation was an important result of combination treatment, ectopic Bcl-xL overexpression led to reversal of GSK-J4 and Belinostat effects in *IDH1*-mutant cells. Moreover, an observed sharp increase in the expression of *DDIT3*/CHOP and its known pro-apoptotic targets *PMAIP1*/NOXA and *BBC3/*PUMA upon combination treatment helped skew the balance from survival under stress to stress-induced apoptosis **(Figure 6J)**. Further validation of this mechanistic model was obtained by significant abrogation of GSK-J4 and/or Belinostat effects in *DDIT3* depleted *IDH1*^*R132H*^ cells.

Finally, the *in vivo* studies with our patient-derived *IDH1*-mutant model support clinical translation of our findings. Only 6-day-treatment of GSK-J4 and Belinostat reduced tumor growth, revealing a therapeutic potential of this combination strategy. Previous work indicates that GSK-J4 is effective on pediatric brainstem glioma *in vivo*, demonstrating its ability to penetrate the blood-brain barrier (BBB)^47^. It was also reported that Belinostat can cross the BBB and has antitumor effects in an orthotopic rat glioma model^48^. Our results add to the growing evidence and list of epigenetic compounds with ability to target brain tumors. However, GSK-J4 is known to have low stability in physiological conditions, and future KDM6 inhibitors with improved metabolic stability and pharmacokinetic properties will need to be developed for clinical translation. In conclusion, our study identifies combined targeting of KDM6A/B and HDACs as a potent and selective therapeutic strategy for the treatment of *IDH1*-mutant glioma.

## ACKNOWLEDGMENTS

Funding was obtained from The Scientific and Technological Research Council of Turkey (TUBITAK) (#2211e) (AK), (219S882) (TBO), Bone Cancer Research Trust (AC, UO), Rosetrees Trust (UO), CRUK (A23900, UO), the LEAN program grant of the Leducq Foundation (UO), the Oxford NIHR Biomedical Research Centre (UO), and the People Programme (Marie Curie Actions) of the European Union’s Seventh Framework Programme (FP7/2007-2013) (UO) under REA grant agreement n° [609305]. The authors acknowledge use of the services and facilities of the Koç University Research Center for Translational Medicine (KUTTAM), funded by the Presidency of Turkey, Presidency of Strategy and Budget.

## AUTHOR CONTRIBUTIONS

Study design: TBO, AK, and UO; data generation: AK, EY, GNS, FS, AC, BI, and SA; data analysis: AK, EY, APC, and TBO; data interpretation: TBO, AK, EY, APC, DPC, IS, HW and UO; supplied reagents: DPC, IS, HW and UO; manuscript draft: AK, EY and TBO; approved final manuscript: all authors.

