## Supplementary Information - Figures S1-S7, Tables S1-S4, Supplementary Methods for "Orthogonal targeting of KDM6A/B and HDACs mediates potent therapeutic effects in *IDH1*-mutant glioma"

### 1. SUPPLEMENTARY FIGURES AND LEGENDS

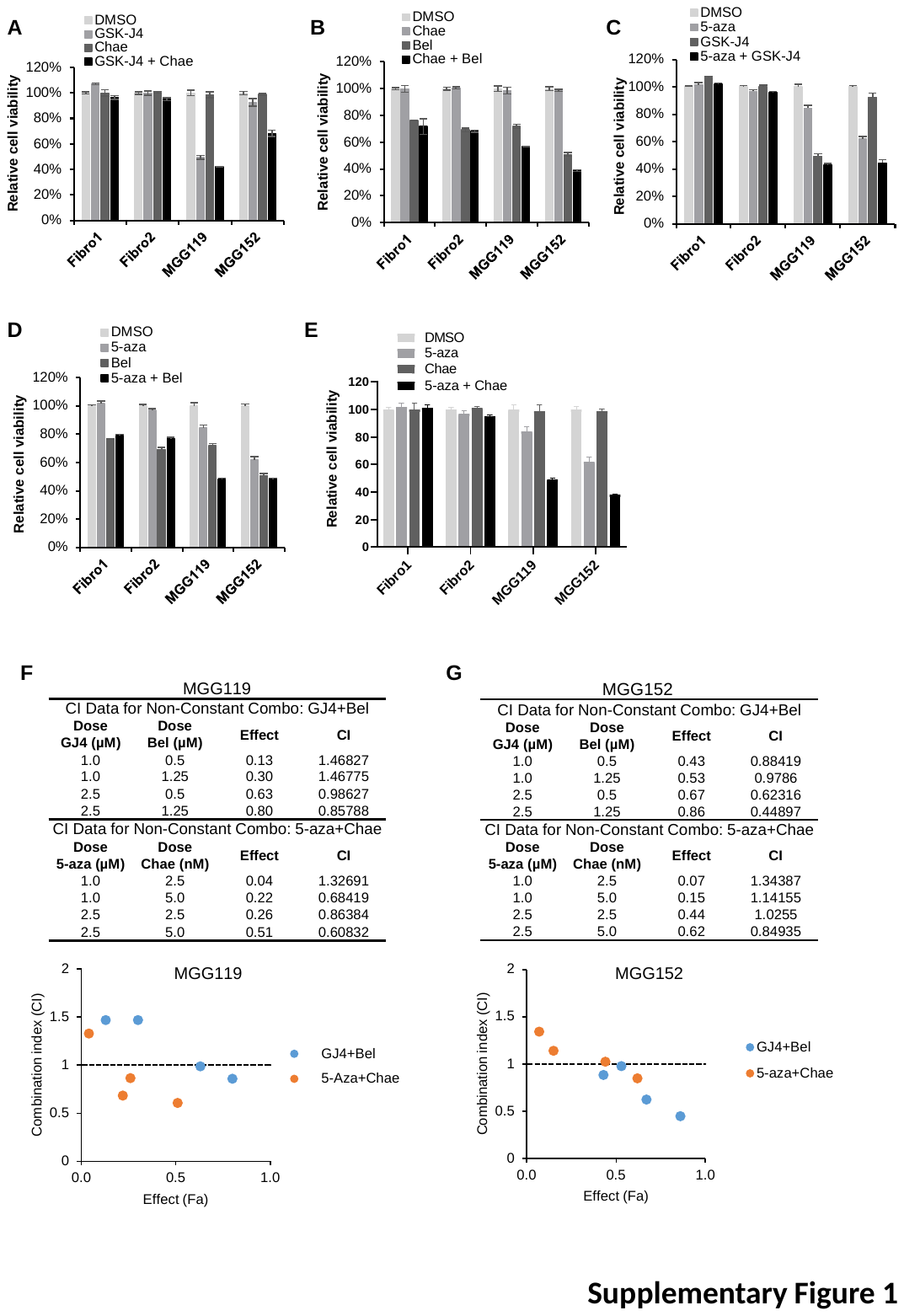

**Supplementary Figure 1. Effect of drug combinations on *IDH1*-mutant cells and combination index calculations.** GBM cells and fibroblasts were treated with dual combinations of screen hits: GSK-J4 and Chatetocin **(A)**, Chaetocin and Belinostat **(B)**, 5-azacytidine and GSK-J4 **(C)**, 5-azacytidine and Belinostat **(D)**, 5-azacytidine and Chaetocin **(E)**. Combination index for GSK-J4 and Belinostat or 5-azacytidine and Chaetocin co-treatments with different doses in MGG119 cells **(F)** and MGG152 cells **(G)** were calculated via CompuSyn software.

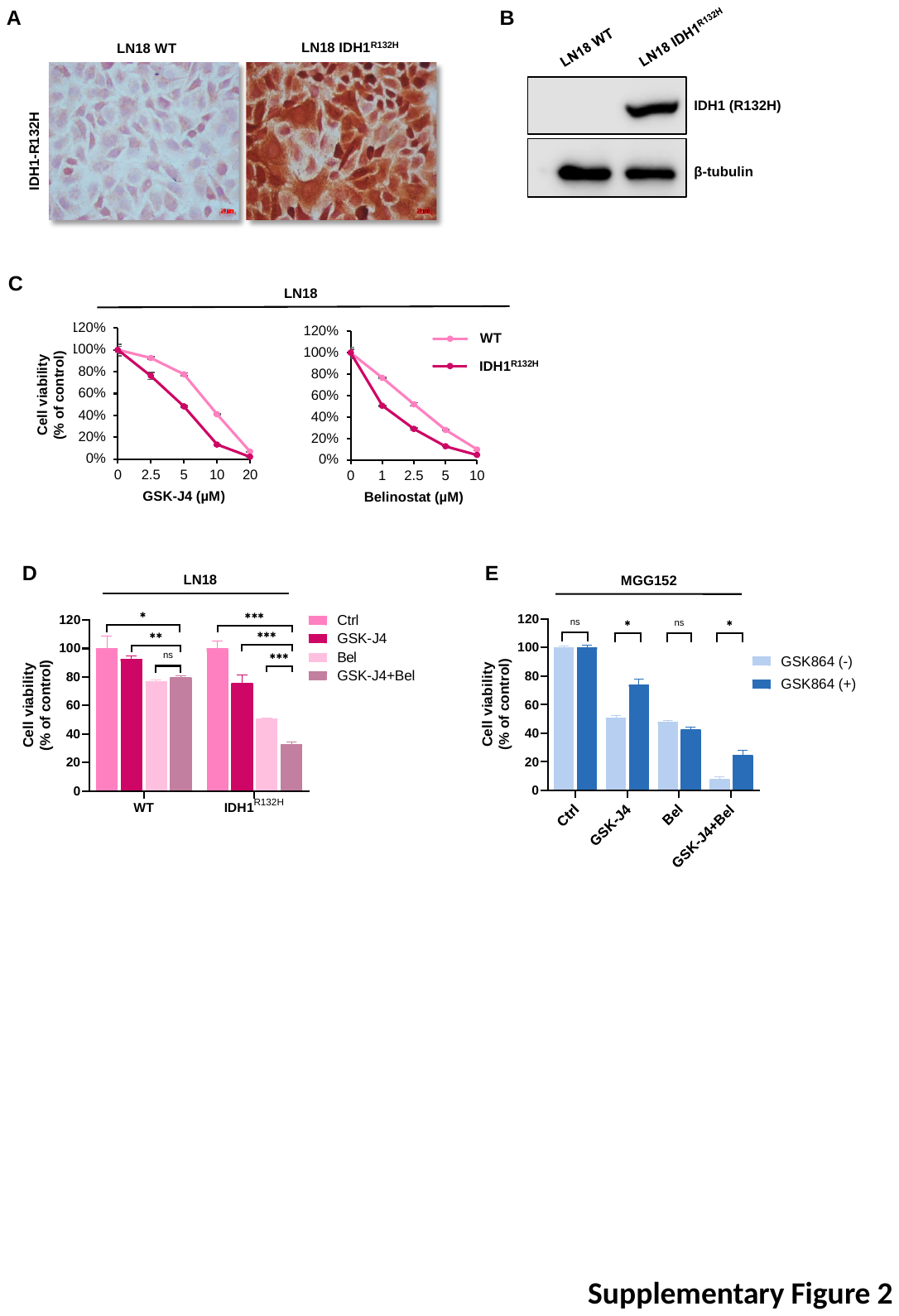

**Supplementary Figure 2. Effects of GSK-J4 and Belinostat combination treatment on LN18 cell line pair.** Validation of mutant IDH1 enzyme (IDH1^R132H^) in LN18 GBM cells via immunohistochemical staining **(A)** and western blot **(B)** using anti-IDH1 (R132H) antibody. **C)** LN18 wild type and *IDH1^R132H^* cells were treated with GSK-J4 or Belinostat individually at different doses for 72h. **D)** LN18 wild type and *IDH1^R132H^* cells were treated with GSK-J4 (2.5 µM) and Belinostat (1 µM) individually or in combination. **E)** Sensitivity of primary MGG152 cells against GSK-J4 and Belinostat treatment was slightly recovered via long-term passaging *in vitro*, in the presence of GSK864. p-values were determined by unpaired t test; ns, non-significant; *p < 0.05; **p < 0.01; ***p < 0.001.

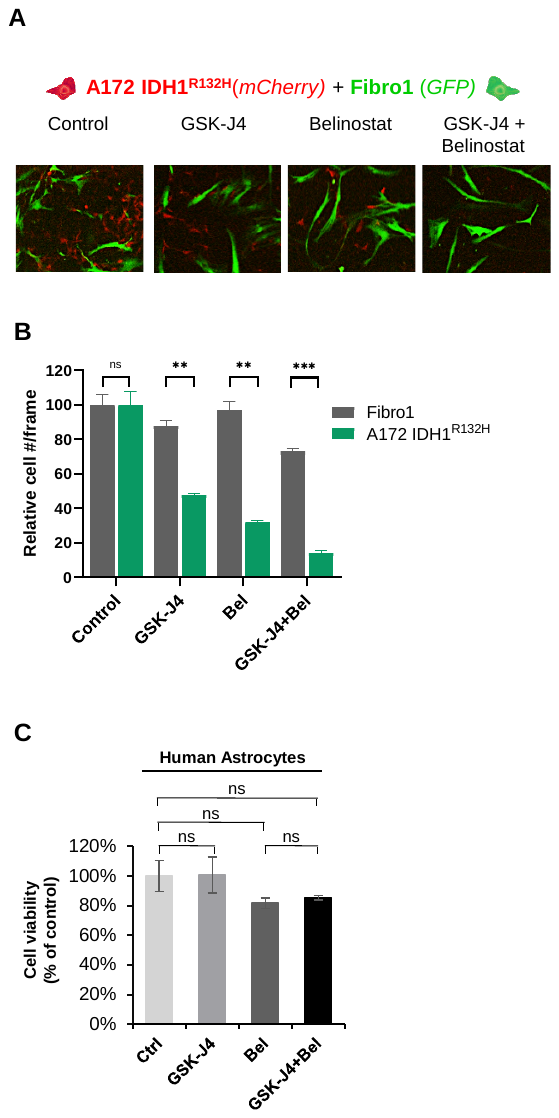

**Supplementary Figure 3. GSK-J4 and Belinostat combination does not affect the viability of non-malignant cells. A)** mCherry-labelled A172 *IDH1^R132H^* cells were co-cultured with GFP-labelled fibroblasts and treated with GSK-J4 and Belinostat individually or in combination. **B)** Quantification of co-culture images after 5 days of treatment with GSK-J4 and/or Belinostat. Images were quantified by ImageJ software. **C)** Viability of normal human astrocytes (NHA) upon combinational treatment of GSK-J4 (2.5 µM) and Belinostat (1 µM). p-values were determined by unpaired t test; ns, non-significant; *p < 0.05; **p < 0.01; ***p < 0.001.

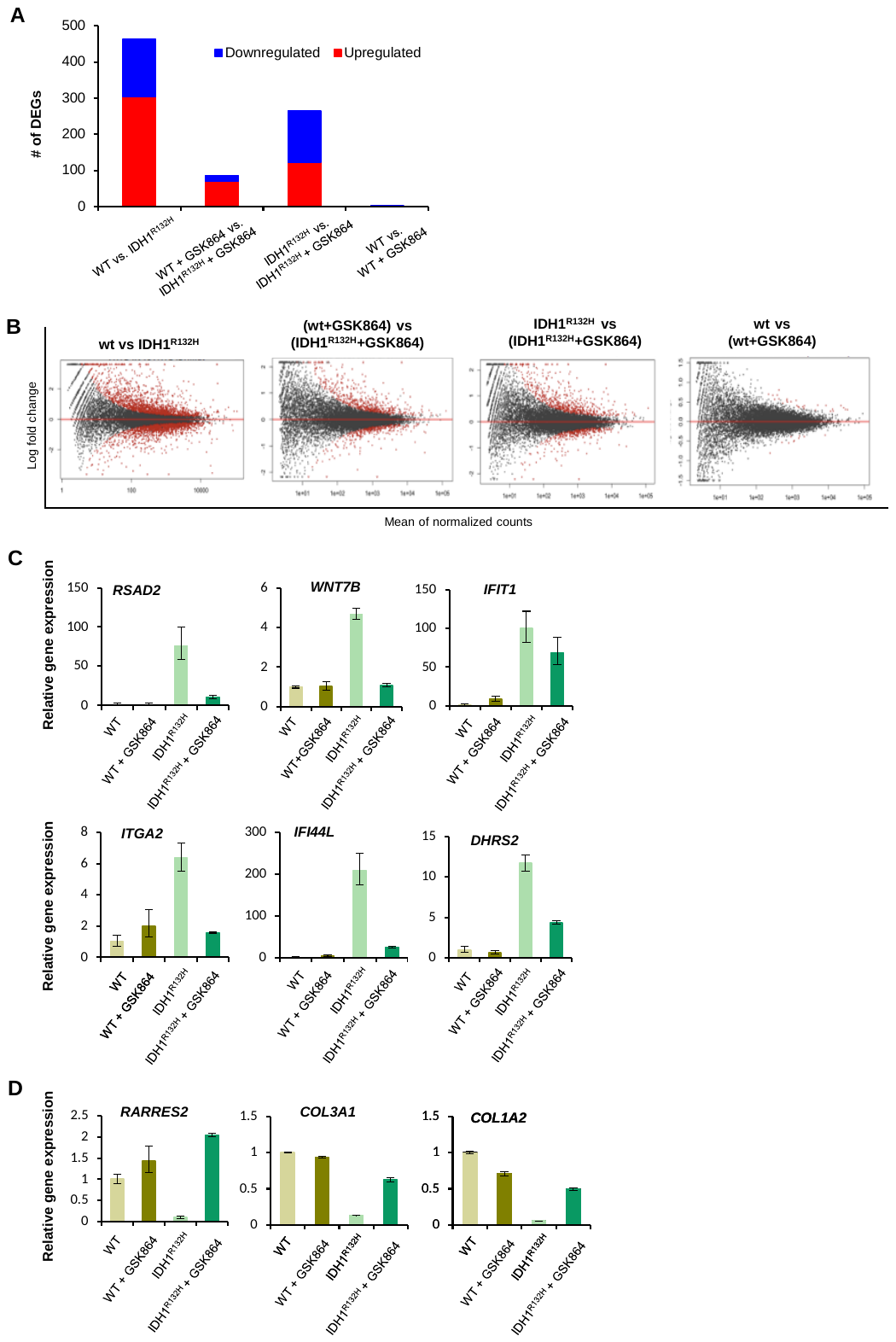

**Supplementary Figure 4. RNA sequencing of A172 WT and *IDH1^R132H^*** **cells. A)** Graph showing numbers of differentially expressed genes (DEGs) in A172 wild type (WT) or *IDH1^R132H^* cells with or without GSK864 treatment. **B)** MA plots showing the DEGs in *IDH1^R132H^* cells in the absence or presence of GSK864. Significantly different counts were shown in red. **C)** qRT-PCR results for validation of DEGs upregulated in *IDH1^R132H^* cells and downregulated back with GSK864 treatment. **D)** qRT-PCR results for validation of DEGs downregulated in *IDH1^R132H^* cells and reactivated with GSK864 treatment.

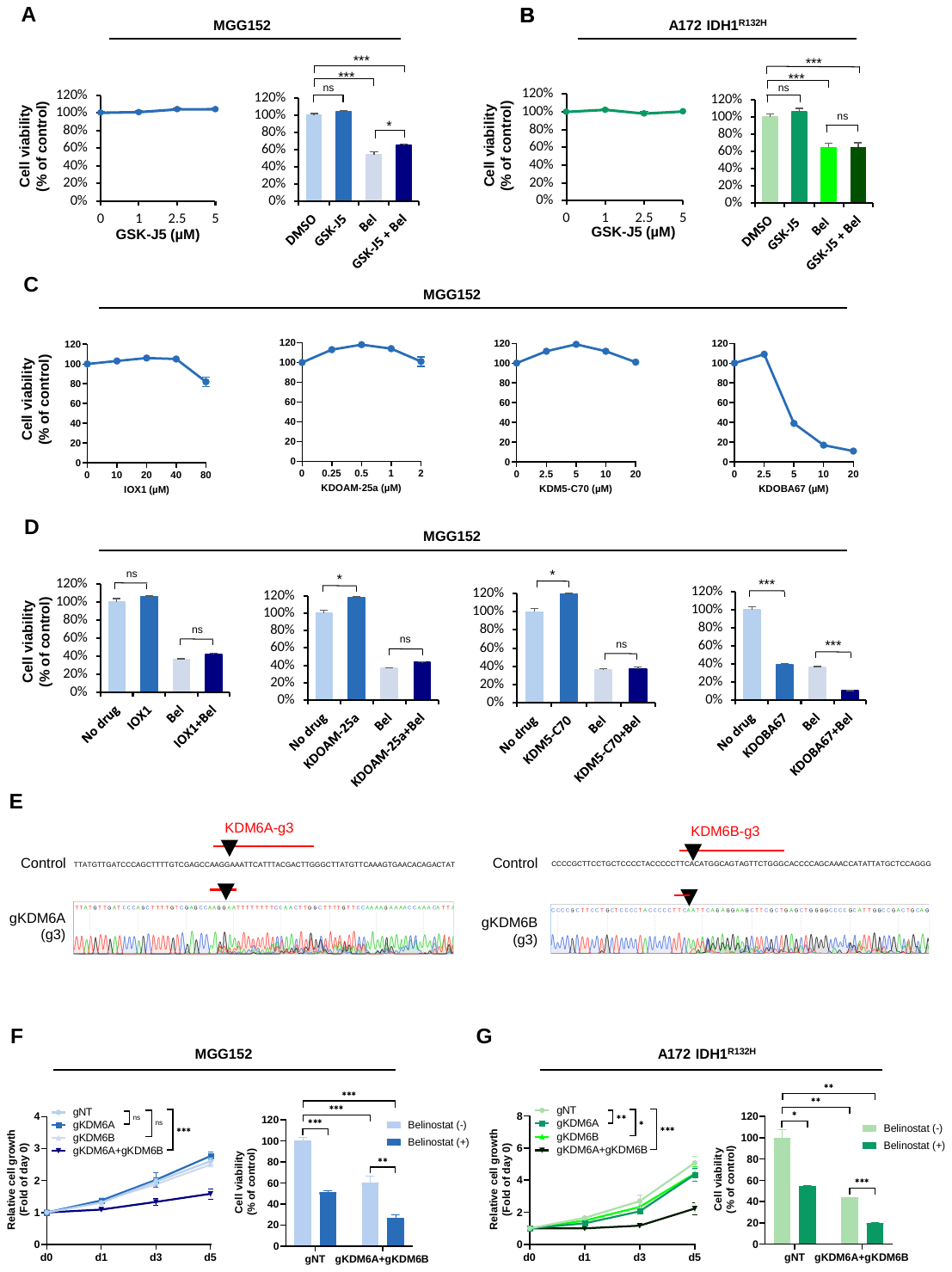

**Supplementary Figure 5. KDM6A and KDM6B inhibition phenocopies the effects observed with GSK-J4 in *IDH1*-mutant cells. A-B)** Viability of primary *IDH1*-mutant MGG152 cells **(A)** or engineered A172 *IDH1^R132H^* cells **(B)**, upon treatment with GSK-J5, an inactive form of GSK-J4, individually or in combination with Belinostat. **C-D)** Viability of primary *IDH1*-mutant MGG152 cells upon treatment with pan 2-OG KDM inhibitor IOX1, KDM5 inhibitors KDOAM-25a and KDM5-C70, or KDM6 inhibitor KDOBA67, which has very similar structure with GSK-J4, individually **(C)** or in combination with Belinostat **(D)**. **E)** Validation of KDM6A and KDM6B knockout by Sanger sequencing. Red bars on the control sequence indicate regions targeted via gRNAs, and black triangles indicate the expected cleavage site. Sanger sequencing reveals frameshifts around the cleavage sites (bottom). **F-G)** Growth of primary MGG152 **(F)** or A172 *IDH1^R132H^* **(G)** cells upon individual or double knockout of *KDM6A* or *KDM6B* compared to non-targeting gRNAs (gNT), and effects of double knockouts on Belinostat treatment in these cells. For growth curves in panel F and G, p-values were determined by 2-way ANOVA test. For all other panels, p-values were determined by unpaired t test; ns, non-significant; *p < 0.05; **p < 0.01; ***p < 0.001.

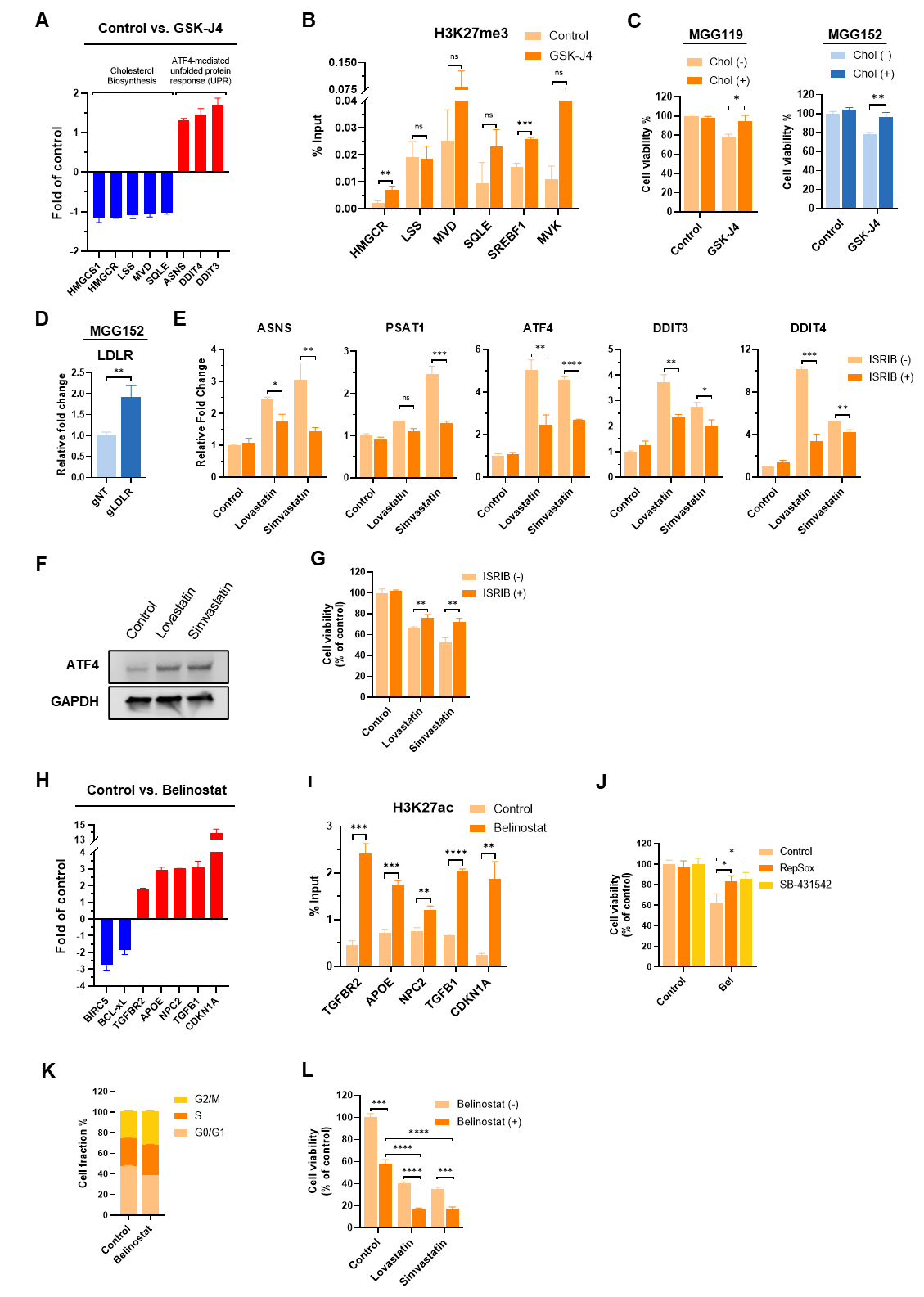

**Supplementary Figure 6. Validation of findings on another primary *IDH1*-mutant MGG119 cells. A)** Expression of genes involved in cholesterol synthesis and ATF4-mediated stress response pathways upon 48h of GSK-J4 treatment. **B)** Relative H3K27me3 level measured by ChIP-qPCR, in the promoter regions of genes involved in cholesterol biosynthesis pathway. **C)** Relative viability of MGG119 and MGG152 cells after 48h of GSK-J4 treatment in the presence or absence of exogenous cholesterol (250 µM). **D)** Relative expression of LDLR gene in MGG152 cells infected with non-targeting (gNT) or LDLR targeting (gLDLR) gRNAs together with dCas9-TET1 and MPH viruses. **E)** Relative expression of genes involved in ATF4-mediated stress pathway upon lovastatin (50 µM) or simvastatin (50 µM) treatment in the presence or absence of ISRIB (1 µM). **F)** Western blot image showing effect of lovastatin or simvastatin treatment on ATF4 levels. **G)** Relative viability of MGG119 cells after 48h of lovastatin or simvastatin treatment in the presence or absence of ISRIB (1 µM). **H)** Expression of genes involved in TGF-β, cholesterol efflux and apoptosis/cell cycle pathways upon Belinostat treatment. **I)** Relative H3K27ac levels measured by ChIP-qPCR, in the promoter regions of genes involved in TGF-β, cholesterol efflux and apoptosis/cell cycle pathways. **J)** Relative viability of MGG119 cells after 48h of Belinostat treatment in the presence or absence of TGF-β inhibitors, RepSox or SB-431542 (0.5 µM). **K)** Graphs representing changes in cell cycle phases after 24h of Belinostat treatment in MGG119 cells. **L)** Relative viability of MGG119 cells after 48h of Lovastatin and Simvastatin treatments in the presence or absence of Belinostat (1 µM). p-values were determined by unpaired t-test; ns, non-significant; *p < 0.05; **p < 0.01; ***p < 0.001, ****p<0.0001.

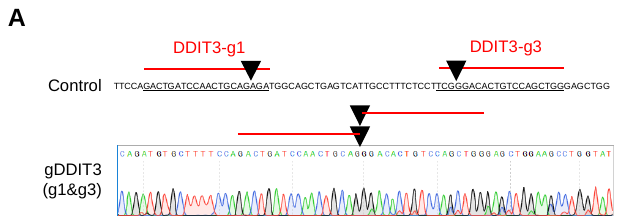

**Supplementary Figure 7. Validation of CRISPR/Cas9 mediated knockout of *DDIT3* gene by Sanger sequencing.** Red bars on the control sequence indicate regions targeted via gRNAs, and black triangles indicate the expected cleavage site. In the bottom parts, Sanger sequencing reveals deletion between the two cleavage sites.

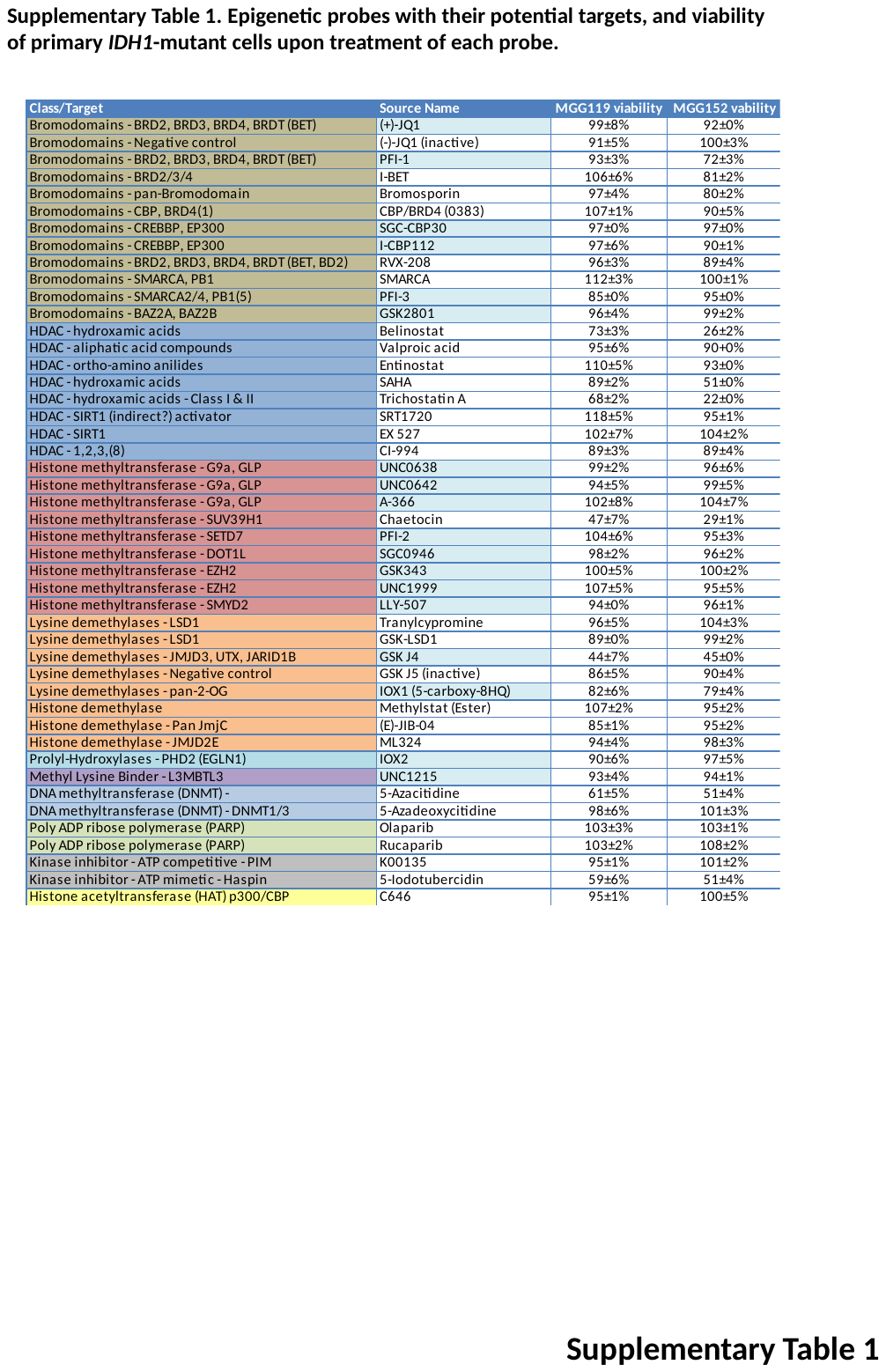

**Supplementary Table 1.** Epigenetic probes in the drug screen with their potential targets, and viability of MGG119 and MGG152 primary *IDH1*-mutant cells upon treatment of each inhibitor.

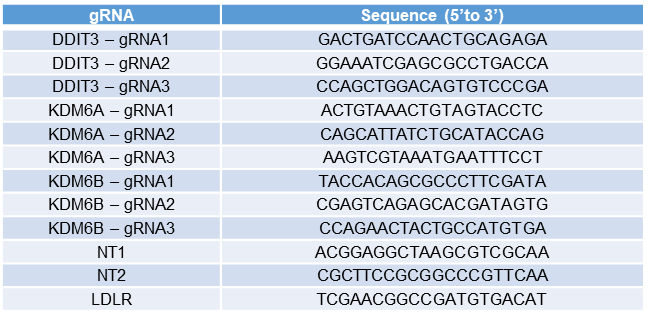

**Supplementary Table 2.** gRNA sequences used for knockout experiments

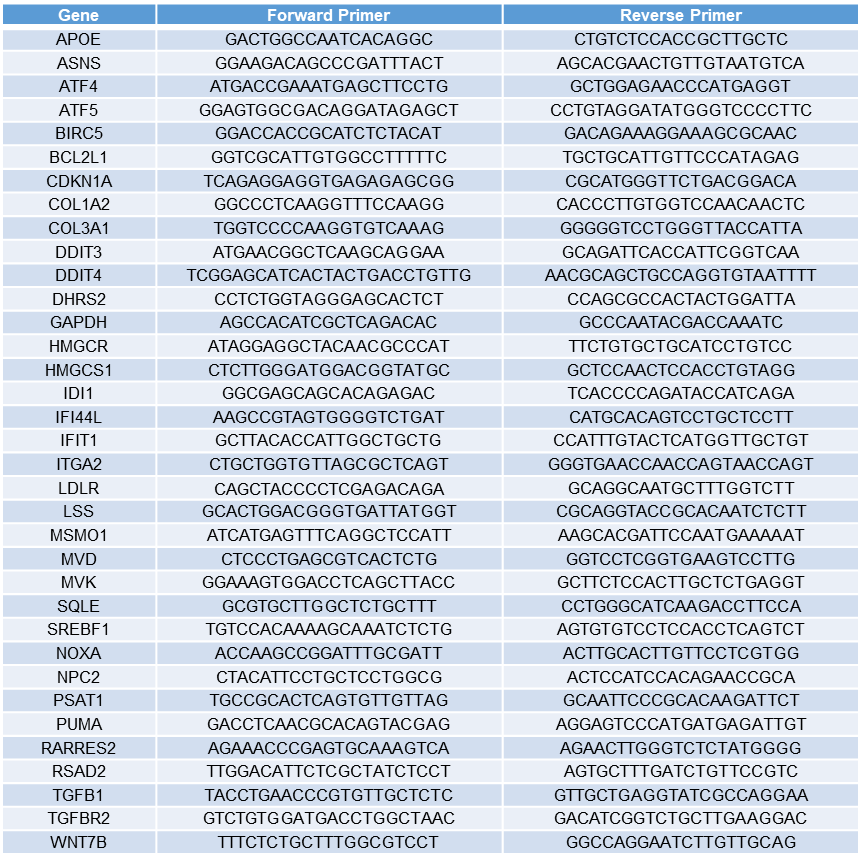

**Supplementary Table 3.** Primer sequences used in qRT-PCR

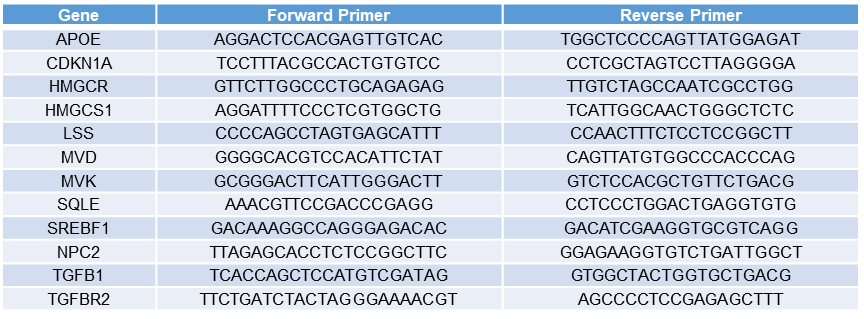

**Supplementary Table 4.** Primer sequences used in ChIP-qPCR

### 2. SUPPLEMENTARY METHODS

### Reagents

The epigenetic chemical probe library was constructed as described^1^ and kindly provided by Prof. Udo Oppermann. Bulk amounts of GSK-J4 and Belinostat were purchased from Adooq Bioscience (CA, USA) for *in vivo* experiments. D-2-Hydroxyglutarate (D2HG) Assay Kit was purchased from Sigma-Aldrich (USA). GSK864 was kindly provided by SGC Toronto. Z-VAD-FMK and Z-FA-FMK were supplied from BD Pharmingen (CA, USA). ISRIB and PERKi were provided by Prof. Udo Oppermann. TMZ was purchased from Selleckchem (USA). Anti-IDH1 R132H (Hu) from mouse (clone: H09) antibody was purchased from Dianova (Germany). Anti-PARP, anti-ATF4 (D4B8), anti-p21 (12D1), anti-H3 (D1H2), anti-H3K27me3 (C36B11), anti-H3K27ac (D5E4), and anti-acetyl-α-tubulin (D20G3) were purchased from Cell Signaling Technology (USA). Anti-GAPDH and anti-α-tubulin were purchased from Abcam (UK). Anti-puromycin (4G11) antibody was purchased from Merck Millipore (USA). GSK-J5, Thapsigargin and Cycloheximide were purchased from Cayman Chemical (USA). D-luciferin was purchased from Biotium (CA, USA). Cholesterol and Simvastatin were purchased from Sigma (USA). Lovastatin was purchased from MedChemExpress (USA).

### Cell culture

293T cells, A172 and LN18 GBM cell lines were purchased from American Type Culture Collection (ATCC, USA). Fibro1 and Fibro2 fibroblast cell lines were established at Koç University School of Medicine from skin biopsies of GBM patients operated at Koç University Hospital (Ethics approval no: 2013-5). Cells were cultured in DMEM (Gibco, USA), with 10% FBS (Gibco, USA) and 1% Pen/Strep (Gibco, USA) in a 37^o^C incubator with 5% CO_2_. Human astrocytes (HAs) were purchased from ScienCell Research Laboratories (CA, USA) and were cultured on Poly-L-Lysine coated plates. MGG119 and MGG152 IDH1 mutant cell lines were established with patient-derived xenograft model^2^. They were cultured as neurospheres in GBM/EF medium consisting of neurobasal medium (Gibco, USA) with 7.5 ml L-Glutamine, 1X B-27 supplement (Gibco, USA), 0.5X N-2 supplement (Gibco, USA), 0.5 ml heparin solution (0.2%, Stemcell Technologies, Canada), 0.5% Pen/Strep, FGF (20 ng/ml), EGF (20 ng/ml).

### Cell Viability and Caspase Activity Assays

Cell viabilities after required treatments were measured via Cell Titer-Glo (CTG) Luminescent Cell Viability Assay (Promega, USA) using a plate reader (BioTek’s Synergy H1, VT, USA). Cells were seeded to clear bottom black side 96-well plates (Corning Costar) as 4000 cells/well and as triplicates for each condition. The next day, they were treated with corresponding chemicals of interest individually or in combination for defined period.

For Caspase 3/7 activity assay, A172 wild type and IDH1 mutant cells were seeded on 96-well plates as 4000 cells/well. The next day, cells were treated with GSK-J4 (2.5 µM) and/or Belinostat (1 µM). After 48h of drug treatments, Caspase 3/7 activities in each well were measured via Caspase Glo 3/7 assay (Promega, USA), according to manufacturer’s instructions.

### Viral packaging and transduction

Retroviral particles from pMIG Bcl-xL (Addgene plasmid #8790), and lentiviral particles from pLenti6.3/TO/V5 containing IDH1^R132H^,^3^ pLentiCRISPRv2 (Addgene #52961), lenti sgRNA(MS2)_zeo (Addgene #61427), lenti MPHv2 (Addgene #89308) and pLEX-dCas9-Tet1 (cloned for unpublished work) were produced in 293T cells as previously described^4^. Adherent GBM cells were seeded as 100.000 cells/well of 6-well plate and transduced with virus containing media and Protamine sulfate (PS, 8 µg/ml). Primary GBM cells cultured as suspension, were transduced with spinfection method (as described in Supplemental Methods). Briefly, neurospheres were resuspended in 2 mL of EF medium supplemented with PS (8 µg/ml) and seeded on 1 well of 6-well plates together with concentrated virus. Then, cells were centrifuged in a plate-centrifuge (Beckman Coulter, USA) at 800g for 90 minutes. Both adherent and suspension cells were incubated overnight in the cell culture incubator, and then, media were refreshed. 48 hours post-transduction, adherent and suspension cells infected with pLentiCRISPRv2 were treated with 2 μg/ml or 1 μg/ml of puromycin, respectively, and selected for 3 days. Suspension cells infected with pLEX-dCas9-TET1, lenti MPHv2 and lenti sgRNA(MS2)_zeo were selected with puromycin (1 μg/ml), hygromycin (75 μg/ml) and zeocin (150 μg/ml), respectively. A172 GBM cells infected with pLenti6.3/TO/V5 containing IDH1^R132H^ were selected with blasticidin (10 μg/ml) for 5-7 days. Efficiency of pMIG BcL-xL infections was monitored via GFP expression under fluorescence microscope.

### CRISPR-mediated knock-out or activation studies

gRNA sequences targeting exon regions of KDM6A and KDM6B genes were obtained from GeCKO library designed by Feng Zhang’s laboratory at Broad Institute^5^. gRNAs targeting DDIT3 gene or LDLR promoter were designed using Chopchop gRNA design tool^6^. All gRNAs synthesized at Macrogen Europe Laboratories as top and bottom strands. They were annealed and cloned into lentiCRISPR v2 plasmid (Addgene #52961) for knockout, and lenti sgRNA(MS2)_zeo plasmid (Addgene #61427) for activation experiments according to protocol published by Zhang Lab^5^. gRNAs targeting each gene were pooled and lentiviruses were produced from these pools for each gene. 2 gRNAs targeting nowhere in human genome were pooled and used as non-targeting control gRNAs (gNT) in further experiments. For LDLR activation, cells were infected with lentiviruses containing dCas9-TET1, MPH and non-targeting or LDLR promoter-targeting gRNAs. Cells were infected with ~MOI of 3 and selected with appropriate antibiotics. Then, growth analysis, viability assays after drug treatments and gene expression analysis were performed. Gene knockouts were validated via Sanger sequencing of targeted regions (**Figure S8**), and activations were validated via qRT-PCR. gRNA sequences were given in **Supplementary Table 3**.

### Total cholesterol assay

Total cholesterol level in A172 and MGG152 cells were measured by Amplex Red Cholesterol Assay Kit (Thermo Fisher, USA) according to the manufacturer’s instructions. Briefly, cell pellets were collected from required conditions. Lipids were extracted in 200 µL of Chloroform: Isopropanol: NP-40 (7:11:0.1) using a sonicator (Biorupter, Diagenode, USA). They were centrifuged at 15.000 x g for 5 min. Supernatants were transferred to new tubes and air dried at 50°C to remove chloroform. Dried lipids were dissolved with 200 µL of Assay Buffer. Standards were prepared by diluting a cholesterol reference to the range of 0 to 20 µM. Reaction buffer was used as negative control, and H_2_O_2_ (10 µM in reaction buffer) was used as positive control. 50 µl from diluted samples, standards and controls were placed on different wells of a 96-well plate. Working solution of Amplex Red reagent containing 2U/ml HRP, 2U/ml cholesterol oxidase and 0.2U/ml cholesterol esterase was prepared and 50 µl of it was added on each sample or control well. Plate was protected from light and incubated at 37 ^o^C for 30 min. Fluorescence levels were measured in a microplate reader (BioTek’s Synergy H1, USA) using excitation detection at 560 nm and emission detection at 590 nm. Cholesterol levels in samples were calculated based on the standard curve obtain by prepared standards.

### Cell cycle analysis

A172 wild type, A172 *IDH1^R132H^* and MGG152 cell pellets were collected after 24h or 48h of either without treatment or treatments with GSK-J4, Belinostat and their combination. They were washed with PBS and fixed with 200 µl of cold 70% ethanol via incubating for 30 min at 4^o^C. Ethanol was added dropwise, and pellets were resuspended either with flicking tube or pipetting gently after each drop. After fixation, cells were centrifuged at 850g for 5 min, and washed 2 times with PBS. To remove RNA content, pellets were resuspended in 50 µl of 100 µg/ml RNase and incubated for 15 min at room temperature. Finally, 200 µl of 50 µg/ml propidium iodide (PI) solution was added on samples, and they were incubated for 30 min at room temperature. Samples were kept in the dark at 4^o^C until analysis was performed. Quantification of PI staining were performed via flow cytometry, and cell cycle intervals were determined by assigning the n peak as G0/G1 phase, n to 2n interval as S phase and 2n peak as G2/M phase.

### YO-PRO-1 staining

A172 wild type and IDH mutant cells were seeded on 24-well plates as 20.000 cells/well. Next day, cells were either kept untreated or treated with GSK-J4 (2.5 µM) and Belinostat (1 µM) in combination. After 48h of drug treatments, media were removed, and cells were kept in fresh media containing 0.1% (v/v) YO-PRO-1 stock solution. Plates were incubated at 37^o^C for 15 min, and fluorescent images were taken directly, as soon as possible upon incubation.

### Western blot and SUnSET assay

Cell pellets were collected after 24h or 48h of indicated drug treatments and stored at -80 ^o^C until the experiments were performed. Pellets were lysed in NP40 lysis buffer (1% NP40, 50 mM Tris buffer pH 7.4, 250 mM NaCl, 5mM EDTA, 50 mM NaF, 0.02% NaN_3_) supplemented with 1X protease inhibitor cocktail set (cOmplete™ ULTRA Tablets, Roche, Germany) and 1mM PMSF prior to usage. Cell lysates were centrifuged at 13.000 rpm for 10 min at 4 ^o^C, and supernatants were collected.

For histone analysis, cell pellets were lysed in Triton Extraction Buffer (TEB) which consists of PBS with 0.5% Triton X 100 (v/v), 2 mM PMSF, 0.02% (w/v) NaN_3_ and 5 mM sodium butyrate, on ice for 10 min. Nuclei were precipitated via centrifuging at 6500 g for 10 min at 4 ^o^C, and washed again with TEB after discarding supernatants. Pellets were resuspended in 0.2N HCl as around 10^7 cells/ml and kept at 4 ^o^C overnight. Next day, lysates were neutralized by adding 1M of NaOH as 1/5 volume of HCl solution. They were centrifuged at 6500 g for 10 min at 4 ^o^C, and supernatants were stored as histone lysates at -20 ^o^C.

For SUnSET assay^7^, MGG152 cells were treated with GSK-J4 and/or Belinostat for 24h or 48h. Thapsigargin (5 µM) or Cycloheximide (5 µM) were used as control treatments. Then, cells were treated with puromycin (10 µM) for 30 min, pellets were collected and lysed in NP40 buffer as explained above.

Protein concentration of lysates from all protocols were determined via Pierce’s BCA protein assay kit (Thermo Fisher Scientific, USA). Lysates were mixed with 4X loading dye which is prepared by mixing 4X Laemmli Sample Buffer (Bio-Rad, USA) with 2-mercaptoethanol in 9:1 ratio, and boiled at 95 ^o^C for 10 min. Appropriate amount of samples and protein ladder (Precision Plus Protein, Bio-Rad, USA) were loaded into gradient SDS polyacrylamide gels (Mini-PROTEAN® TGX™ Precast Gels, Bio-Rad, USA), and run at 25 mA for 40 min. Then, protein transfer was performed via Trans-Blot® Turbo™ RTA Mini PVDF Transfer Kit (Bio-Rad, USA) with manufacturer’s defined transfer protocol. Membrane was blocked with 5% non-fat dry milk for 1h with gentle shaking at room temperature. Then, blocking buffer was replaced with primary antibody diluted in PBST with 2% BSA and 0.02% NaN_3_ by gently shaking overnight at 4 ^o^C. Next day, antibody solution was removed, and membrane was washed 3 times with PBST for 15 min each. Then, it was incubated with secondary antibody diluted in 5% milk solution for 1h at RT and washed 3 times with PBST for 15 min each. Membrane was incubated with Pierce™ ECL Western Blotting Substrate (Thermo Fisher Scientific, USA) for 5 min at dark, and visualized by Odyssey ® Fc Imaging System (LI-COR Biosciences, USA).

### RNA sequencing

All samples were studied as duplicate. Total RNAs were extracted with Nucleospin RNA kit (Macherey-Nagel, Germany), according to manufacturer’s protocol, and stored at -80 ^o^C until sending for RNA-seq analysis. The RNA-seq libraries were prepared using NEBNext Ultra Directional RNA library prep kit for Illumina (NEB, USA), according to manufacturer’s instructions. Prior to the analysis, 42 bp paired-end reads were quality controlled using FastQC (version 0.11.4)^8^, aligned using HISAT2 (version 2.1.0)^9^, and assigned to annotated features using featureCounts (subread version 1.5.0-p2). The Fasta and GTF files for the human genome were obtained from the Ensembl FTP site release 75. Computational pipelines were used by calling scripts from the CGAT toolkit to analyze the next generation sequencing data (<https://github.com/cgat-developers>)^10^. Data have been deposited in NCBI's Gene Expression Omnibus (with the accession number of GSE134120). Differentially expressed genes (DEGs) were identified using DESeq2^11^. Analysis of enriched pathways were performed using Gene Set Enrichment Analysis (GSEA) and eXploring Genomic Relations (XGR) softwares^12,13^.

### qRT-PCR experiments

Cell pellets were collected after specified treatments and stored at -80 ^o^C until the following experiments were performed. Total RNAs were isolated via Nucleospin RNA kit (Macherey-Nagel, Germany), according to manufacturer’s instructions, and stored at -80 ^o^C. cDNAs were produced from 1000 ng of total RNA for each condition. qRT-PCR was performed by using SYBR green mix (Roche, Switzerland), and Lightcycler 480 instrument (Roche, Switzerland). Relative gene expressions were calculated by using ΔΔCt method, and GAPDH as reference gene. Primers used in qRT-PCR experiments were given in **Supplementary Table 2.**

### ChIP-qPCR experiments

Pellets were collected from MGG119 and MGG152 cells after 48h of GSK-J4 or Belinostat treatment. For crosslinking, 2x10^6^ cells for each condition were dissolved in 1% formaldehyde (in PBS), and shaken at room temperature for 10 min. Glycine with the final concentration of 0.125 M was added to solution to stop crosslinking, and tubes were shaken at room temperature for 5 min. Then, pellets were precipitated by centrifuge at 1500 rpm for 5 min, and washed with cold PBS for 2 times. After the second wash, pellets were dissolved in 100 µl of ChIP lysis buffer containing 1% SDS, 10 mM EDTA, 50 mM Tris-HCl (pH=8.1) and protease inhibitors (0.2 mM PMSF, 1 µg/ml Aprotinin, 1 µg/ml Leupeptin) and taken into 1.5 ml-sonication tubes. Samples were sonicated in Biorupter (Diagenode, USA) with refrigerated water (4 ^o^C) for 3 x 12 cycles. Between each cycle, 30 seconds break were set, and between two 15-cycles tubes were pipetted up and down, and put into the sonicator back. After sonication, cell debris were removed by centrifuge at 17500g for 10 min at 10 ^o^C. Supernatants were transferred to new tubes and diluted 10 times by ChIP dilution buffer containing 0.01% SDS, 1.1% Triton-X 100, 1.2 mM EDTA,16.7 mM Tris-HCl (pH=8.1), 167 mM NaCl. 10% of diluted samples were collected as input control, and decrosslinked with the addition of NaCl to the final concentration of 0.2 M, and incubation overnight at 65 ^o^C. The next day, samples were taken and stored at -20 ^o^C until further use. Rest of the samples were used for immuno-precipitation by addition of 5 µg from antibody of interest and shaking at cold room overnight. The next day, 108 µl of A/G Sepharose beads (slurry) were added to 900 µl of suspensions containing immuno-complexes. Samples were shaken at cold room for 2h. Beads were collected by placing sample tubes on a magnetic holder. Supernatants were removed by aspiration and washed sequentially by low salt wash buffer containing 0.1% SDS, 1% Triton-X 100, 2 mM EDTA, 20 mM Tris-Hcl (pH=8.1), and 150 mM NaCl, high salt wash buffer containing 0.1% SDS, 1% Triton-X 100, 2 mM EDTA, 20 mM Tris-Hcl (pH=8.1), and 500 mM NaCl, LiCl wash buffer containing 0.25 M LiCl, 1% IGEPAL-CA 630, 1% deoxycholic acid, 1 mM EDTA, 10 mM Tris (pH=8.1), and two times with Tris-EDTA (TE). For each wash, beads were dissolved in 1 ml of buffer, shaken for 5 min at cold room, placed on magnetic holder and supernatants were aspirated. After last wash, beads were dissolved in 150 µl of elution buffer containing 1% SDS and 100 µM of sodium bicarbonate in dH_2_O and shaken for 15 min at room temperature. After beads were collected with magnetic holder, supernatants were transferred to a new tube. Beads were dissolved in a fresh 150 µl of elution buffer. Both supernatants were combined, and beads were discarded. Samples were decrosslinked as described above for the inputs. Then, both input and immunoprecipitated samples were purified with Qiagen PCR purification kit, according to the manufacturer’s instructions. Purified DNAs were used as template for the qPCR. Levels of investigated histone marks were calculated as input % for each condition and compared with each other.
